# Sub-millisecond conformational transitions of short single-stranded DNA lattices by photon correlation single-molecule FRET

**DOI:** 10.1101/2021.04.09.439239

**Authors:** Brett Israels, Claire S. Albrecht, Anson Dang, Megan Barney, Peter H. von Hippel, Andrew H. Marcus

## Abstract

Thermally-driven conformational fluctuations (or ‘breathing’) of DNA plays important roles in the function and regulation of the ‘macromolecular machinery of genome expression.’ Fluctuations in double-stranded (ds) DNA are involved in the transient exposure of pathways to protein binding sites within the DNA framework, leading to the binding of regulatory proteins to single-stranded (ss) DNA templates. These interactions often require that the ssDNA sequences, as well as the proteins involved, assume transient conformations critical for successful binding. Here we use microsecond-resolved single-molecule Förster Resonance Energy Transfer (smFRET) experiments to investigate the backbone fluctuations of short oligothymidine [oligo(dT)*_n_*] templates within DNA constructs that can also serve as models for ss-dsDNA junctions. Such junctions, as well as the attached ssDNA sequences, are involved in the binding of ssDNA binding (ssb) proteins that control and integrate the mechanisms of DNA replication complexes. We have used these data to determine multi-order time-correlation functions (TCFs) and probability distribution functions (PDFs) that characterize the kinetic and thermodynamic behavior of the system. We find that the oligo(dT)_*n*_ tails of ss-dsDNA constructs inter-convert, on sub-millisecond time-scales, between three macrostates with distinctly different end-to-end distances. These are: (*i*) a ‘compact’ macrostate that represents the dominant species at equilibrium; (*ii*) a ‘partially extended’ macrostate that exists as a minority species; and (*iii*) a ‘highly extended’ macrostate that is present in trace amounts. We propose a model for ssDNA secondary structure that advances our understanding of how spontaneously formed nucleic acid conformations may facilitate the activities of ssDNA associating proteins.

**Significance Statement:** The genetic information of living organisms is encoded as sequences of nucleic acid bases in DNA, and is protected by the thermodynamically stable secondary structure of the Watson-Crick double helix. The processing and manipulation of gene sequences by ‘macromolecular machines’ requires that stable segments of duplex DNA be disrupted, and that single-stranded (ss) DNA templates be transiently exposed to the binding sites of DNA associating proteins within the cellular environment. Here we elucidate some of the defining features that control the stability and dynamics of ssDNA secondary structure, using time-resolved methods to detect the presence of transient unstable conformations. Understanding the nature of these instabilities is central to elucidating the mechanisms by which ssDNA templates facilitate protein binding and function.

## 1. Introduction

The genome of living cells exists primarily as B-form double-stranded (ds) DNA, which is packaged in the cell nucleus as chromatin (in eukaryotes) or nucleoid bodies (in bacteria) (1). During various stages of the cell cycle, segments of single-stranded (ss) DNA are exposed by the action of the DNA replication helicase to make available the ssDNA templates required by the DNA polymerases for the synthesis of the complementary daughter strands for cell division, and to allow access of other functional and regulatory proteins to these transiently formed ssDNA sequences. Within the cell, and in functional complexes such as the replisome, these exposed ssDNA segments are coated by ssDNA binding (ssb) proteins that function to protect the genetic information from nucleases and to favorably configure the single strands for further manipulation by the ‘machinery of genome expression’ (2, 3). These proteins interact with ssDNA to form stable, cooperatively bound nucleoprotein filaments, in which the bound ssDNA chain segments are nearly fully extended to their maximum contour lengths permitted by the sugar-phosphate backbones, resulting also in largely unstacking the attached bases (4, 5). Because such fully extended conformations of ssDNA are likely to be unstable in the absence of bound protein, a fundamental question arises concerning how ssb protein-ssDNA interactions evolve along a molecular reaction coordinate to form extended nucleoprotein filaments.

Unlike the cooperatively-structured and stabilized dsDNA conformation, many possible conformations can be adopted by ssDNA in solution, and the specifics of these conformations depend on a variety of interactions between the nucleic acid components with their aqueous surrounds. These interactions include base stacking and unstacking, intra-chain repulsion between adjacent backbone phosphates, counterion condensation along the polyelectrolyte ssDNA backbone, binding and orientation of polar water molecules around these counterions and the backbone phosphates, strain within the sugar-phosphate backbone and increases in configurational entropy of the backbone as a consequence of base-unstacking and the concomitant rotation around multiple single-bonds of the extended backbone (6–8).

The cumulative effect of these interactions is a free energy landscape that determines the probability with which a given ssDNA configuration will occur. In general, the coordinate space over which the free energy landscape is defined can be partitioned into distinct sub-spaces, herein referred to as ‘macrostates,’ which correspond to local basins of stability separated by relatively high activation barriers (9). Each macrostate consists of a ‘family’ of energetically-degenerate microscopic conformations (or ‘microstates’) that rapidly interconvert (on tens-of-nanoseconds time-scales) in a non-Markovian manner (10–12). In contrast, higher thermal barriers separate conformations associated with different macrostates, such that their interconversion occurs on tens-of-microseconds and longer time scales. The relatively high activation barriers associated with macrostate interconversion ensures that the observed dynamics of the overall system can be accurately described using a kinetic master equation in which the rate constants contain no ‘hidden variables’ and memory effects need not be considered (13).

In this work, we present microsecond-resolved single-molecule Förster resonance energy transfer (smFRET) experiments to monitor the conformational fluctuations of short single-stranded oligo-deoxythymidine [oligo(dT)_*n*_] templates of varying length (*n* = 14 and 15) and polarity (3’ versus 5’). We note that these constructs have been used in previous studies as model systems to monitor the dynamics of the non-cooperative and cooperative binding and assembly of the T4 bacteriophage ssb proteins (gene product 32, or gp32) onto target ssDNA lattices (14, 15). The binding site size of the gp32 monomer is 7 nucleotides (nts) so that for the templates with a total length, *n*, of 14 nts there is excess room for the non-cooperative binding of a single gp32 monomer or, alternatively, precisely enough room for the cooperative assembly of a gp32 dimer. For templates with *n* = 15 nts, there are two possible conformations for the gp32 dimer-bound state; either at nucleotide positions 1 – 14 or 2 – 15. Furthermore, the gp32 proteins are expected to assemble onto ssDNA lattices in a polar manner, so that the assembly mechanism will depend on the polarity of the ss-ds DNA junction. Our previous studies showed that gp32 proteins bind to, and dissociate from, the ss-dsDNA junction as monomers on tens-of-milliseconds time scales, and that gp32 monomer-bound intermediates must encounter (also, on tens-of-milliseconds time scales) a ‘productive’ ssDNA conformation near the ss-dsDNA junction to permit a second ssb protein to form a cooperatively-bound gp32 dimer-cluster (14). By examining the sub-millisecond dynamics of the ssDNA lattice fluctuations in these constructs in the absence of proteins, we may ask whether short-lived conformational intermediates – which can serve as potential transition states for ssb binding – play a role in the assembly mechanism.

Each of the oligo(dT)_*n*_ conformations that we investigated were formed within the overhanging ‘tail’ of a ss-dsDNA junction construct, in which the distal end of the ssDNA chain is labeled with the FRET donor chromophore Cy3 and the conjugate DNA strand is labeled, at the ss-dsDNA junction, with the FRET acceptor chromophore Cy5 (see Fig. 1). Such ss-dsDNA constructs labeled in this way were previously used by Lohman and Ha and co-workers in related studies with the E. coli SSB system using standard smFRET methods (3, 16, 17). Our experiments detect individual fluorescence photons from a continuously irradiated single molecule sample using instrumental methods (18) and data analyses developed previously (14, 19). Because these experiments resolve individual photon detection events, the dominant contribution to measurement uncertainty is the sparse statistical sampling of the ensemble-averaged signal rate (20), which varies stochastically in time due to the conformational fluctuations of the macromolecule. The time-dependent signal rates of the Cy3 donor (*D*) and the Cy5 acceptor (*A*) FRET-pair chromophores are given by *I*_*D*(*A*)_(*t*) = *N*_*D*(*A*)_(*t*)/*T_w_*, where *N*_*D*(*A*)_ is the number of photons detected during a fixed time window *T_w_* (= 10 *μ*sec) beginning at the time *t*. Thus, the smFRET efficiency, *E_FRET_*(*t*) = *I_A_*(*t*)/[*I_D_*(*t*) + *I_A_*(*t*)], is a time-averaged signal, which represents an average of the macromolecular configurations detected during the sampling period *T_w_*.

**Figure 1.**
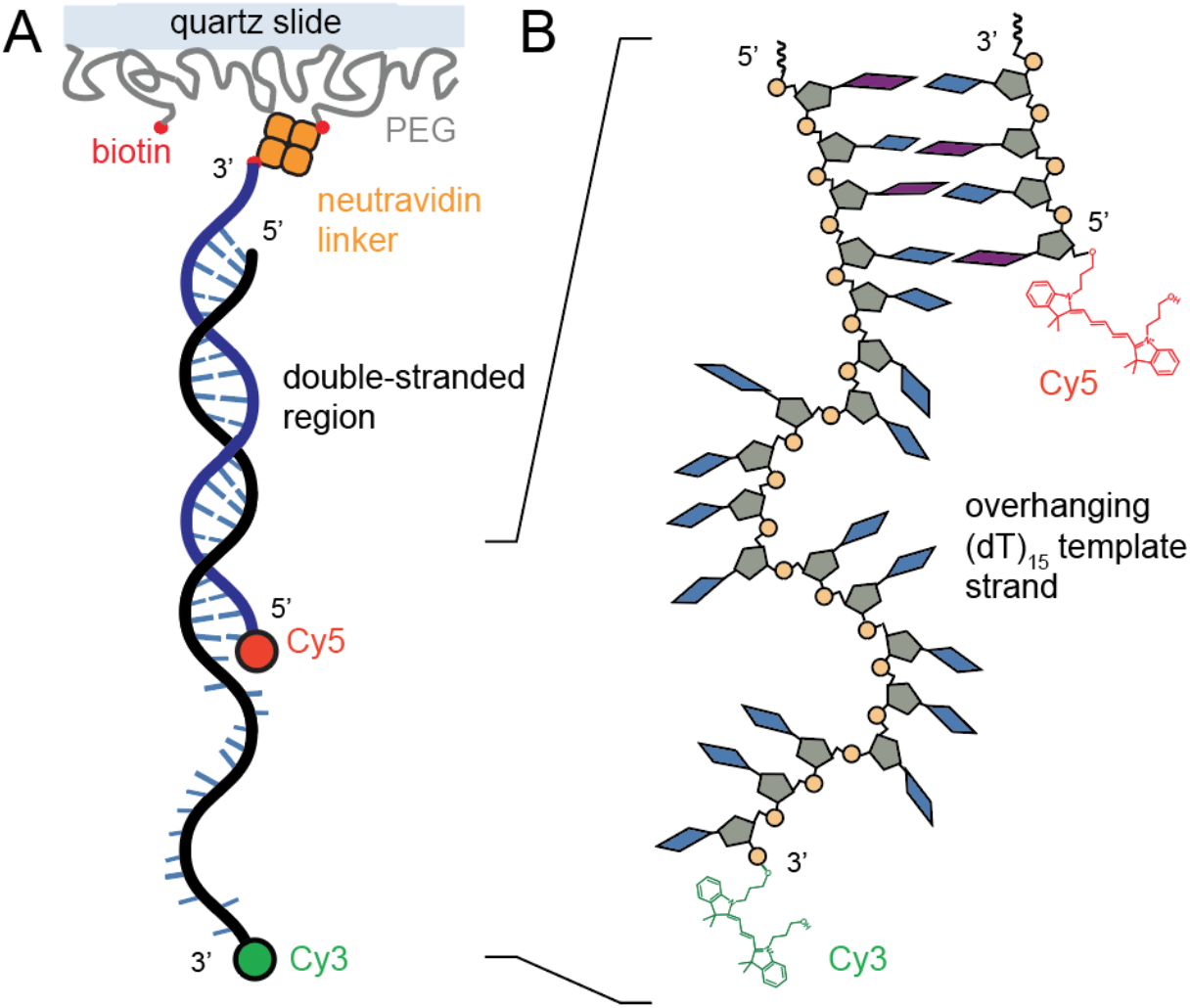
Schematic of one of the four single-stranded (ss) – double-stranded (ds) DNA constructs [3’-Cy3/Cy5-oligo(dT)_15_-ss-dsDNA – see Table 1] used in our experiments. (***A***) The double-stranded region of the ss-dsDNA construct was attached to the surface of a fused silica quartz (SiO_2_) slide coated withPEG and containing biotin-neutravidin linkers. (***B***) The Cy3 donor chromophore was attached to the distal end of the oligo(dT)_*n*_ overhanging ‘tail’ region of the ss-dsDNA construct, and the Cy5 acceptor was attached to the opposite strand at the ss-dsDNA junction, as shown.

Our microsecond-resolved smFRET measurements contain both structural and kinetic information about the microscopic configurations of the oligo(dT)_*n*_ templates. To extract the maximum amount of information possible from our data, we applied an ‘inverse-problem’ procedure for analyzing photon correlation measurements of this type (14, 19). From the photon count data stream, we determined the following three ensemble-averaged functions of the smFRET signal: (*i*) the time-averaged probability distribution function (PDF) of the mean FRET efficiency *P*(*E_FRET_*); (*ii*) the two-point time correlation function (TCF), 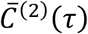, which describes the average correlations between two successive measurements separated by the time interval *τ*; and (*iii*) the four-point TCF, 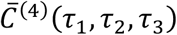, which describes the average correlations between four successive measurements separated by the time intervals *τ*_1_, *τ*_2_ and *τ*_3_. The above functions can be used to characterize the equilibrium and kinetic properties of a multi-step conformational transition pathway at equilibrium. While the two-point TCF, 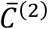, contains information about the characteristic relaxation times of the system, the four-point TCF, 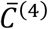, contains additional information about the exchange times between pathway intermediates (19, 21–23).

We simulated the PDF and the TCFs using a master equation (24, 25) that models the system as a network of coupled macrostates, each of which is distinguished by a different mean FRET efficiency. Kinetic rate constants were varied as input parameters to the master equation to achieve the ‘best fit’ between simulated and experimentally-derived functions. As we discuss further below, we compared the results of a linear versus cyclical three-state network model from which we obtained optimized values for the forward and backward rate constants that connect the conformational macrostates, in addition to the probability of observing each macrostate. The optimized rate constants so obtained determine the free energy barriers that separate each macrostate (applying the Arrhenius equation), and the optimized probabilities determine the free energy minima of each macrostate (applying the Boltzmann distribution).

A major finding of this work is that all four of the oligo(dT)_*n*_ templates that we investigated undergo conformational dynamics over a broad range of time scales, spanning tens-of-microseconds to hundreds-of-milliseconds. We observed that all four of these systems interconvert between three distinct macrostates: (*i*) a ‘compact’ and relatively stable macrostate; (*ii*) a ‘partially extended’ macrostate of intermediate stability; and (*iii*) a ‘highly extended’ macrostate of relatively low stability. The free energy minima that we determined for the three macrostates increase rapidly with decreasing FRET efficiency, suggesting that the primary mechanism of macrostate destabilization is the loss of configurational entropy associated with increasing chain extension (10). Moreover, our results indicate the existence of transition state barriers that are intrinsic to the oligo(dT)_*n*_ templates themselves, and which prolong the lifetimes of extended chain configurations relative to the more stable compact configurations. The existence of metastable extended chain conformations suggests that these short-lived species may serve as intermediates along the reaction pathway for the formation of ssb nucleoprotein clusters or filaments.

## 2. Materials and Methods

### 2.1 Single-Stranded (ss) – Double-Stranded (ds) DNA Constructs

We performed microsecond-resolved smFRET experiments on four ss-dsDNA constructs with a ssDNA oligo(dT)_*n*_ overhanging ‘tail’ region that varied in length (*n* = 14 and 15) and polarity (3’ and 5’, see Table 1). In Fig. 1 we show a representative schematic diagram using the 3’-Cy3/Cy5-oligo(dT)_15_-ss-dsDNA construct. These samples were fluorescently labeled using the Cy3/Cy5 FRET donor-acceptor chromophore pair, as indicated.

**Table 1.**
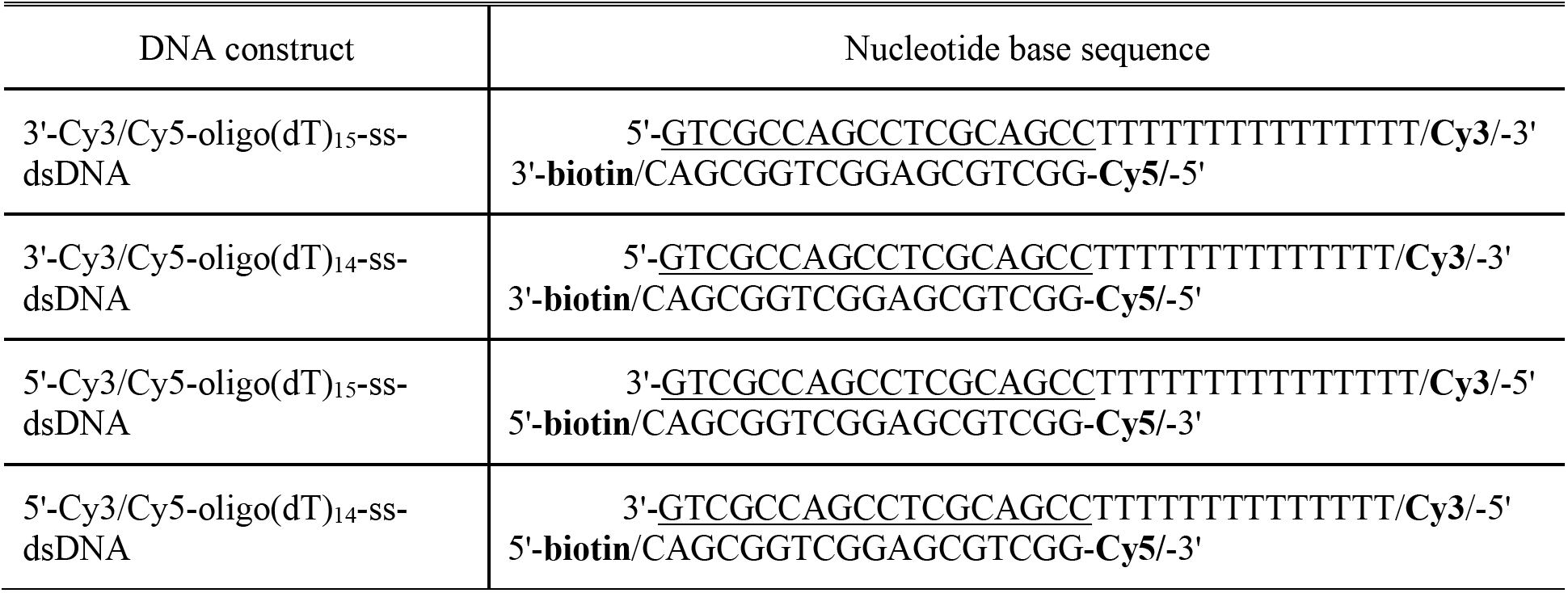
Nucleotide base sequences for ss-dsDNA constructs studied in this work.

### 2.2 Sample Preparation

Sample solutions were prepared in an aqueous buffer containing 100 mM NaCl, 6 mM MgCl_2_ and 10 mM Tris (pH 8.0), as described previously (14, 18, 26). Oxygen scavenging and triplet quenching reagents were used to extend the excited singlet state lifetime and to minimize triplet state blinking (27, 28). Microfluidic sample chambers were constructed using fused silica quartz microscope slides, as shown in Fig. S1*A* (29). Single molecules were attached to the slide surface using a neutravidin linker, which binds strongly to biotin attached to both the ss-dsDNA constructs (Table 1) and to the poly(ethylene glycol) (PEG) layer that coats the slide surface (see Fig. 1*A*). Additional details of the sample preparation are included in the SI.

### 2.3 Microsecond-Resolved Single-Molecule (sm)FRET

We performed our smFRET experiments on the ss-dsDNA constructs using instrumentation and procedures described previously (14, 18). Additional details of the data acquisition and signal calibration are described in the SI. A schematic diagram of the instrumental setup is shown in Fig. S1*B*. The sample was illuminated using a continuous wave laser at 532 nm set up in a prism-type total internal reflection fluorescence (TIRF) configuration. The laser beam was focused to a 50 μm diameter spot at the sample and the incident power was adjusted to 10 mW. The Cy3 donor (*D*) and Cy5 acceptor (*A*) fluorescence from a single ss-dsDNA construct was imaged through a pinhole, and spatially separated into *D* and *A* paths using a dichroic beam splitter. Individual photons from the *D* and *A* paths were detected using avalanche photodiodes (APDs), and each detection event was assigned a ‘time-stamp’ with 0.1 μs resolution and recorded to the computer hard disk. In post-data acquisition, we determined the time-dependent *D*(*A*) signal rates according to *I*_*D*(*A*)_(*t*) = *N*_*D*(*A*)_(*t*)/*T_w_*, where *N*_*D*(*A*)_ was the number of *D*(*A*) photons detected during an adjustable dwell period *T_w_* (≥ 10 *μ*sec) at time *t*. The mean total fluorescence photon detection rate, 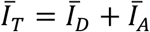, was typically ~15,000 – 20,000 cps. The signal ‘background rate’ was determined to be ~ 2,000 cps by scanning the *xy*-stage position away from the center of a molecule. Thus, the signal-to-background ratio of our experiments is ~ 10. Each single-molecule data set was recorded for a total duration of thirty seconds. From the time-dependent signal rates we determined the FRET efficiency according to *E_FRET_*(*t*) = *I_A_*(*t*)/*I_T_*(*t*).

## 3. Theoretical Background

In our experiments, we monitored the oligo(dT)_*n*_ end-to-end distance *R_DA_* between the *D* (Cy3) and *A* (Cy5) chromophore labels of the ss-dsDNA constructs (see Fig. 1) through measurements of the smFRET efficiency: *E_FRET_* = [1 + (*R_DA_*/*R*_0_)^6^]^−1^. Here *R*_0_ = 0.211(*J_DA_k*^2^*n*^−4^Φ_*D*_)^1/6^ is the characteristic Förster distance (in units of Å) where Φ_*D*_ is the *D* fluorescence quantum yield, *n* is the refractive index of the medium, *J_DA_* is the overlap integral between the *D* emission and the *A* absorbance spectra, and *κ*^2^ is the transition dipole-dipole orientation factor (30). The orientation factor 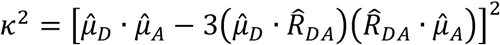 accounts for the relative angles between the *D* and *A* chromophores where 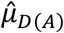 is the unit vector that points in the direction of the *D*(*A*) transition dipole moment and 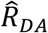 is the unit vector that points from the *D* to the *A*. For cases in which an isotropic distribution of relative *D* and *A* orientations can be assumed, which would require that at least one of the two dyes be freely rotating around its ‘tether,’ the orientation factor takes on the value *κ*^2^ = 2/3 and the Förster distance *R*_0_ ≅ 56 Å (31, 32). In the remainder of this work, we focus on the properties of the observable *E_FRET_* as an indicator of the end-to-end distance, *R_ee_* (assumed ≈ |*R_DA_*|), of the oligo(dT)_*n*_ ‘tail’ regions of the ss-dsDNA constructs that we studied, although we return to the interpretation of the observable in terms of *R_ee_* in the Conclusions.

### 3.1 Macrostates and Free Energy Minima

The oligo(dT)_*n*_ templates can adopt a broad range of microscopic chain configurations (i.e., microstates). Assuming there are *N* total microstates available at a given temperature *T*, the probability that the *i*th microstate occurs is given by the Boltzmann distribution (33)

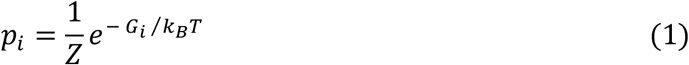

where *k_B_* is Boltzmann’s constant, *G_i_* is the Gibbs free energy of the *i*th microstate, and 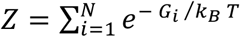 is the partition function. We note that the normalized probabilities sum to unity: 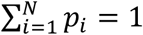. Individual microstates are expected to interconvert on tens-of-nanosecond time scales, which is much faster than the microsecond resolution of our experiments (10). Thus, the ‘observed’ *D* and *A* fluorescence signals each represents a time-average over the possible microstates that occur at a particular time *t* during the sampling period *T_w_* (= 10 *μ*sec). It is useful to define a ‘macrostate’ as a collection of pseudo-degenerate microstates that occupy a common, locally stable region of the free energy landscape. We may then regard each *D* and *A* fluorescence photon detection event as a stochastic sample of the partial-ensemble-average of microstates defined within the phase-space boundary of a single macrostate.

In our analysis, we model the probability distribution function (PDF) of *E_FRET_* values as a sum of contributions from *M* macrostates 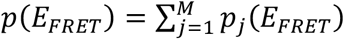, where the *j*th macrostate has the Gaussian form 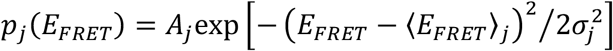, with mean value 〈*E_FRET_*〉_*j*_, standard deviation *σ_j_*, and amplitude *A_j_*. The Gaussian model for the distribution of configurational microstates (within a given macrostate) is consistent with a random-flight, finite-length polymer chain (10, 34). Macrostates that contain a relatively large number of pseudo-degenerate microstates contribute a correspondingly broad distribution of *E_FRET_* values. Thus, the relative probability that the *j*th macrostate is present at equilibrium is given by the integrated area of its component Gaussian 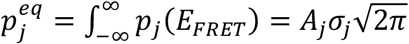. We note that the sum of normalized probabilities over all macrostates is equal to unity: 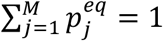. Applying Eq. (1) to macrostates, we may thus write the free energy of the *j*th macrostate (in units of *k_B_T*) as a function of *E_FRET_*

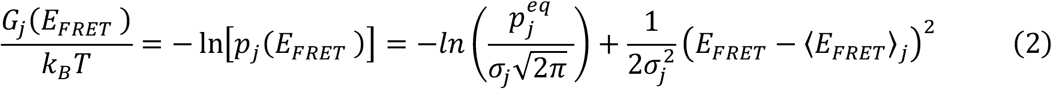

In the context of the above model, the free energy of the *j*th macrostate is represented as a parabolic function of *E_FRET_* with vertical displacement 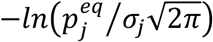. The total free energy surface as a function of *E_FRET_* can then be constructed by summing over the individual macrostate contributions according to 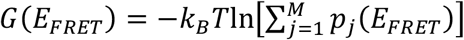.

### 3.2 Free Energies of Activation

We assume that under equilibrium conditions, the microstates of the oligo(dT)_*n*_ template are partitioned amongst *M* quasi-stable macrostates, and there exist conformational reaction channels that permit discreet, stochastic transitions to occur from the *i*th to the *j*th macrostate. In addition, we make the standard approximation that these transformations can be modeled as continuous processes (24, 35). We thus define the time-dependent population of the *i*th macrostate, *p_i_*(*t*), as a continuous, single-valued function that obeys the coupled, first-order ordinary differential equations 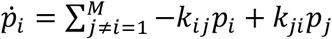. Here the forward rate constant *k_ij_* is associated with the transition from macrostate *i* to macrostate *j*, and the backward rate constant *k_ji_* is associated with the transition from macrostate *j* to macrostate *i*. The rate constant *k_ij_* is related to the free energy of activation 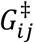, which is given by the Arrhenius equation (33)

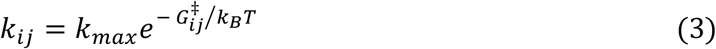

In Eq. (3), the maximum rate constant *k_max_* corresponds to a barrierless process, to which all the thermally activated processes of the system are referenced.

### 3.3 Two-Point and Four-Point Time-Correlation Functions (TCFs)

We obtain kinetic information from our time-resolved smFRET data through our analyses of two-point and four-point time-correlation functions (TCFs) (19). We define the time-dependent fluctuation of the *E_FRET_* observable *δE_FRET_*(*t*) = *E_FRET_*(*t*) – 〈*E_FRET_*〉, where 〈*E_FRET_*〉 is the mean value determined from a time series of measurements performed on a single molecule. The two-point TCF is the average product of two successive measurements separated by the interval *τ*

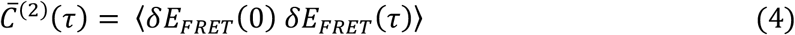

In Eq. (4), the angle brackets indicate a running average over all possible initial measurement times. As we discuss further below, the function 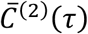 contains information about the minimum number of possible macrostates of the system, each macrostate’s mean FRET efficiency value, in addition to the characteristic time scales of state-to-state interconversion. The two-point TCF may be expressed using a standard statistical mechanical model according to:

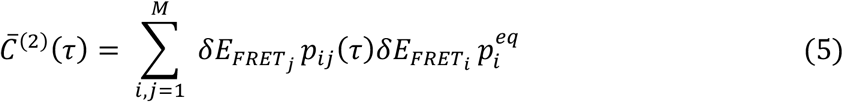

Here 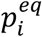 is the probability that the *i*th macrostate is present at equilibrium, 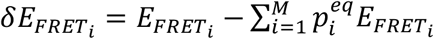 is the value of the fluctuation observable corresponding to the *i*th macrostate, and *P_ij_*(*τ*) is the conditional probability that the system undergoes a transformation from macrostate *i* to macrostate *j* during the interval *τ*. Equation (5) states that 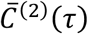 is the statistically weighted average product of consecutive (two-point) observations, which are expected to occur within the interval *τ* as the system undergoes spontaneous transitions between the *M* macrostates.

We define the four-point TCF as the time-averaged product of four successive measurements separated by the intervals *τ*_1_, *τ*_2_, and *τ*_3_, respectively.

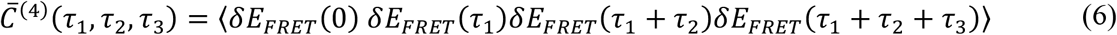

The function 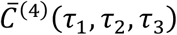 contains information about the roles of intermediates within the conformational kinetic pathways that affect the rates of interconversion between macrostates. The first term on the right-hand-side of Eq. (6) may be modeled according to:

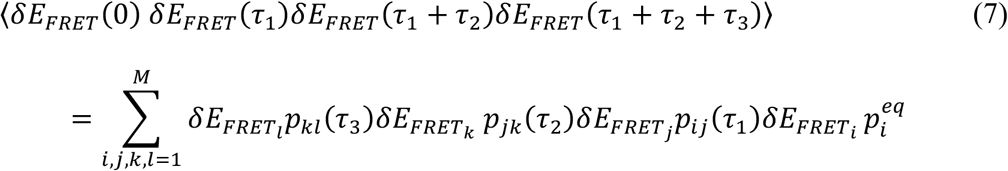

Equation (7) expresses 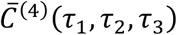 in terms of the weighted average of four-point products of the observable *δE_FRET_*, in which a transition occurs from macrostate *i* to macrostate *j* during the interval *τ*_1_, a subsequent transition occurs from macrostate *j* to macrostate *k* during the period *τ*_2_, and a final transition occurs from macrostate *k* to macrostate *l* during the interval *τ*_3_. We may thus view the four-point TCF as the *τ*_1_, *τ*_2_, *τ*_3_-dependent weighted average of four-point transitions (i.e., elementary kinetic pathways) that connect the *M* macrostates. As we discuss in the next section, the two-point and four-point TCFs described by Eqs. (5) and (7), respectively, in addition to the equilibrium PDF can be modeled theoretically using a kinetic master equation.

### 3.4 Master Equation Analysis

In order to model the ensemble-averaged quantities discussed in the previous sections, it is necessary to determine for a given conformational reaction scheme the equilibrium distribution of macrostates, 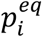, and the time-dependent conditional probability, *p_ij_*(*τ*), that the system will undergo a transformation from macrostate *i* to macrostate *j* during the interval *τ*. These probabilities are solutions to the master equation, which is the set of coupled first-order ordinary differential equations that describe the time-dependent macrostate populations.

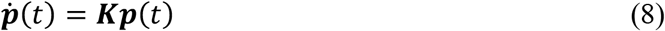

In Eq. (8), ***p***(*t*) = [*p*_1_(*t*), *p*_2_(*t*),…, *p_M_*(*t*)] is the time-dependent macrostate population vector, and 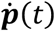 is its time-derivative. The *M* × *M* rate matrix *K* contains the rate constants *k_ij_* according to

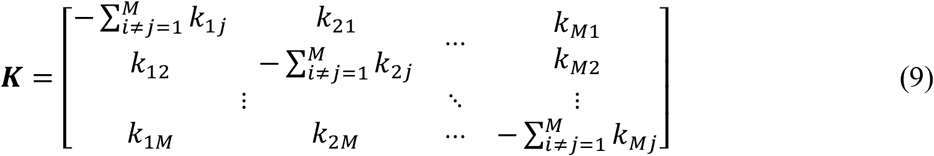

The rate matrix ***K*** is written in the ‘site-basis’ of elementary kinetic steps that comprise the coupled reaction scheme. The off-diagonal elements *K_ij_* = *k_ij_* correspond to the gain in population of macrostate *j* due to the transition *i* → *j*. For cases in which there is no transition *i* → *j, K_ij_* = 0. The diagonal elements are written 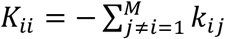, and represent the sum of all reactions that deplete population from macrostate *i*. Equation (8) must satisfy completeness, 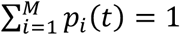, and the detailed balance conditions 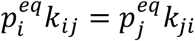 with 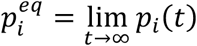. Moreover, the vector of equilibrium macrostate populations ***p**^eq^* can be obtained by solving the master equation subject to the time-stationary boundary condition 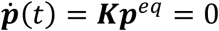.

Given a particular set of rate constants, we solve Eq. (8) by finding the eigenvalues, *λ*_1_, *λ*_2_,… *λ_M_*, and the eigenvectors, ***v***_1_, ***v***_2_,…, ***v_M_***, which satisfy the eigenvector equation ***Kv**_i_* = *λ_i_**v**_i_*. The eigenvalues are equal to the characteristic relaxation rates of the coupled dynamical system, and the eigenvectors represent the collective modes (i.e., linear combinations of elementary transitions) associated with the characteristic relaxations. An important symmetry property of the rate matrix ***K*** is that the elements within each of its columns sum to zero (i.e., 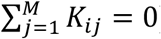), which results in *λ*_1_ = 0 and the remaining eigenvalues *λ*_2_,… *λ_M_* < 0 (36).

A general solution to the master equation [Eq. (8)] is

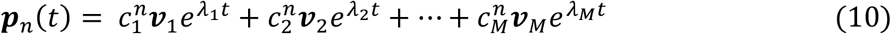

where the constants 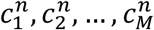 are expansion coefficients that depend on the specific initial condition, which we indicate by the index *n* (25). Equation (10) can be further simplified by making the substitution *λ*_1_ = 0 and taking the infinite-time limit, so that 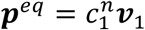 for all possible initial conditions.

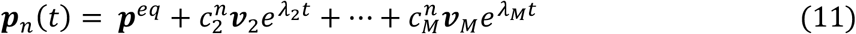

The conditional probabilities needed to evaluate the two-point and four-point TCFs [Eqs. (5) and (7), respectively] are obtained by constraining Eq. (11) using appropriate boundary conditions. The conditional probability *p_ij_*(*t*) is the probability that at time *t*, the population vector has unit occupancy of the *j*th macrostate, given that at time zero there was unit occupancy of the *i*th macrostate. Thus, for example, by applying the initial condition *n* = *i* to Eq. (11) with *p_i_* (0) = *1* and *p*_*j*≠*i*_ (0) = 0, we may determine the expansion coefficients 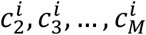. The conditional probability *p_ij_*(*t*) is then equal to the *j*th element of the population vector given by Eq. (11), according to

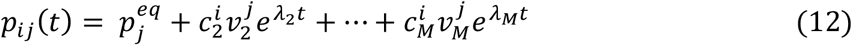

In principle, the above procedure can be used to determine the *p_ij_*(*t*)s analytically for all of *M*^2^ possible combinations of *i,j* ∈ {1, 2,…, *M*}, provided the number of states *M* is relatively small (19). In practice, the *p_ij_*(*t*)s are determined numerically by writing Eq. (11) in the eigenbasis of the ***K*** matrix and carrying out the similarity transformation ***p**_n_*(*t*) = ***U***[***e^λt^***]***U***^−1^***p**_n_* (0), where the unitary matrix ***U*** = [***v*_1_, *v*_2_,…, *v_M_***] and the matrix [***e^λt^***] is diagonal with non-zero elements [*e^λt^*]_*ii*_ = *e^λ_i_t^* (24). Additional details of this procedure are given in the SI.

We note that in the infinite-time limit Eq. (12) recovers the expected equilibrium population values, 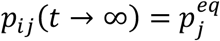. Furthermore, the conditional probabilities *p_ij_*(*t*) are each a sum of *M* − 1 decaying exponential functions of time, with decay constants equal to the eigenvalues of the ***K*** matrix. Substitution of Eq. (12) into Eq. (5) shows that the two-point TCF is also a sum of *M* – 1 exponential terms: 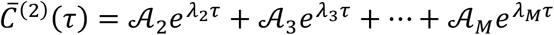, where the 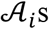 are the relative weights of the collective relaxation processes. Similarly, the four-point TCF can be shown to be a sum of (*M* – 1)^2^ terms: 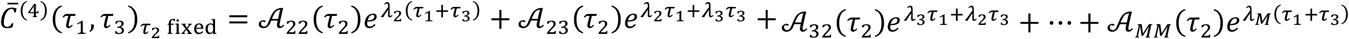 (19). Thus, the number of exponential terms that contribute to the two-point and four-point TCFs indicate the minimum number of macrostates *M* that underlie the observed dynamics.

## 4. Results and Discussion

For each of the four ss-dsDNA constructs that we studied (see Table 1), we recorded time series of single-molecule photon data streams, each for a duration of thirty seconds (see Sect. 2.3). The results of our analyses indicate that all four of the ss-dsDNA constructs exhibit very similar behaviors, as one might expect when focusing on conformational changes within ssDNA sequences only, although the four constructs behave differently at slower time scales in the presence of ssb (gp32) proteins (14, 15, 37). In the following discussion, which deals only with ssDNA conformational changes, we focus on the results of one of the ss-dsDNA constructs [3’-oligo(dT)_15_]. We summarize our results for all four of the ss-dsDNA constructs that we studied in the SI. These additional data sets will serve as reference states in our subsequent considerations of ssDNA-ssb interactions (37).

We begin by making some qualitative observations. In Fig. 2, we show example trajectories of the donor *D* (Cy3, green) and acceptor *A* (Cy5, red) fluorescence intensities constructed from our raw single-molecule photon correlation measurements of the 3’-Cy3/Cy5-oligo(dT)_15_-ss-dsDNA construct. The fluorescence intensities are calculated according to *I*_*D*(*A*)_(*t*) = *N*_*D*(*A*)_(*t*)/*T_w_*, where *N*_*D*(*A*)_(*t*) is the number of *D*(*A*) photons recorded within a finite sampling window *T_w_* beginning at time *t*. An additional example trajectory is shown in Fig. S2 of the SI. We see that the *D* and *A* fluorescence intensities fluctuate in an anti-correlated manner due to the varying inter-chromophore FRET coupling, and indicates that these signals depend on conformational changes of the 3’-oligo(dT)_15_ tail region of the ss-dsDNA construct.

**Figure 2.**
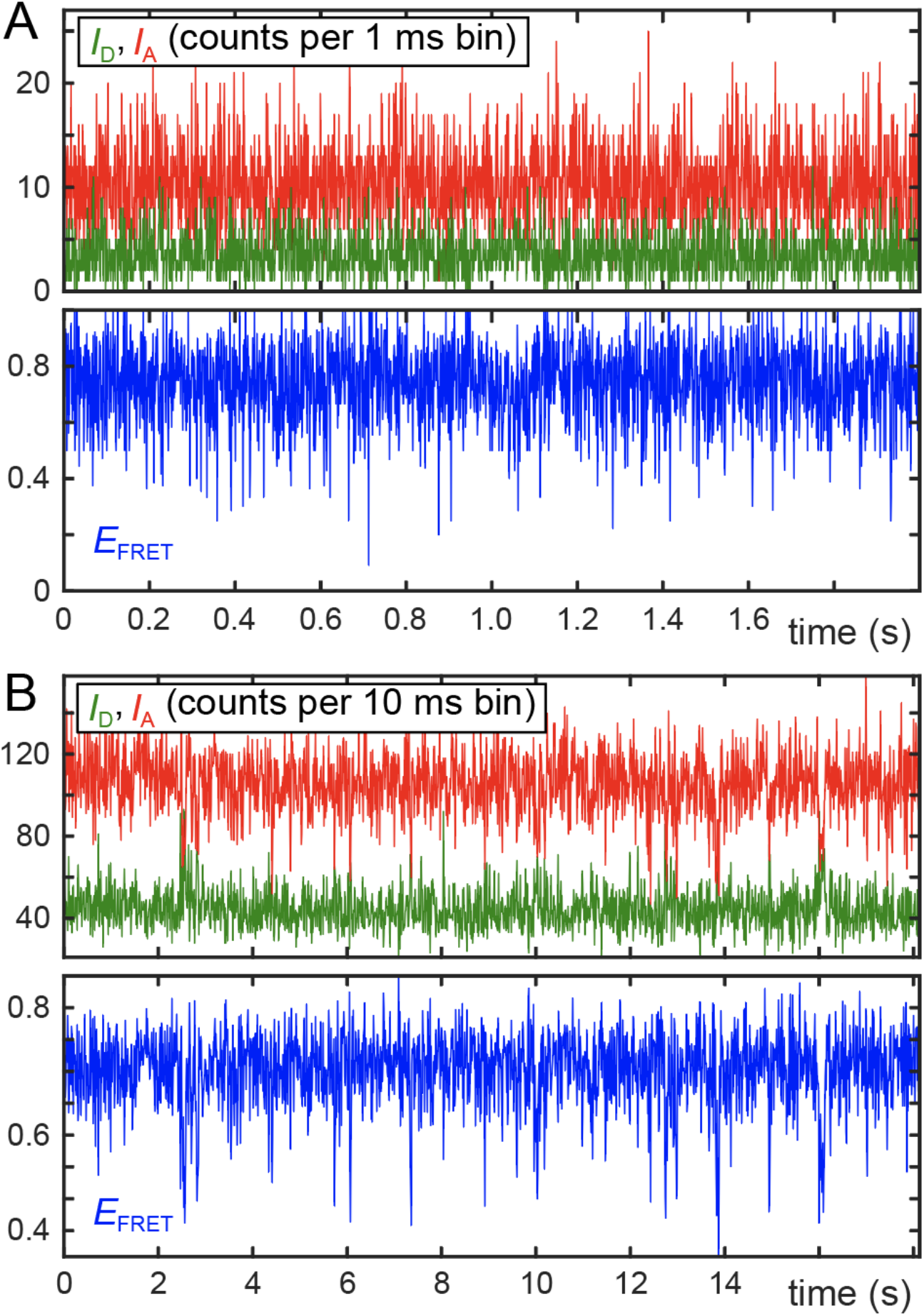
Example smFRET trajectories for the 3’-Cy3/Cy5-oligo(dT)_15_-ss-dsDNA construct. The donor *D* (Cy3, green) and acceptor *A* (Cy5, red) fluorescence intensities, *I*_*D*(*A*)_(*t*) = *N*_*D*(*A*)_(*t*)/*T_w_*, are shown in **panel *A*** with sampling window resolution *T_w_* = 1 ms over a duration of ~ 2 seconds. The FRET efficiency *E_FRET_*(*t*) is plotted in the bottom panel (blue). In **panel *B*** are shown the same data using bin resolution *T_w_* = 10 ms over a duration of ~ 18 seconds. The average Signal-to-Noise Ratio (SNR) per sampling window is 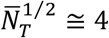 for *T_w_* = 1 ms, and 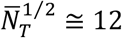 for *T_w_* = 10 ms.

The *D* and *A* intensities are used to calculated the FRET efficiency, according to *E_FRET_*(*t*) = *I_A_*(*t*)/[*I_D_*(*t*) + *I_A_*(*t*)] (blue). We show these trajectories for two different values of the sampling window: *T_w_* = 1 ms (Fig. 2*A*) and *T_w_* = 10 ms (Fig. 2*B*). For this particular photon data stream, the mean total signal flux was 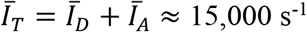. We note that the trajectory appears ‘noisier’ for the case of the *T_w_* = 1 ms sampling window in comparison to the *T_w_* = 10 ms window (see also Fig. S2). This is due to the measurement uncertainty associated with the statistical sampling of photons during the finite sampling window *T_w_* at a constant low-level flux. An average Signal-to-Noise Ratio (SNR) can be assigned to the integrated signals shown in Fig. 2, 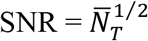, where 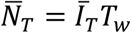 is the mean number of detected photons per sampling window. Thus, for *T_w_* = 1 ms the average 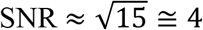, and for *T_w_* = 10 ms the average 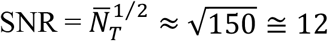. Upon further inspection of the signal trajectories shown in Fig. 2*B*, we see that the majority of FRET efficiency values are clustered near *E_FRET_* ~ 0.7. Moreover, we observe that infrequent transitions occur to short-lived states with FRET efficiency values centered near *E_FRET_* ~ 0.5, and even less frequent transitions occur to short-lived states with *E_FRET_* < 0.5. In the analysis that follows, we quantify this behavior using statistical functions.

We analyzed our raw photon correlation measurements by constructing probability distribution functions (PDFs), in addition to two-point and four-point time-correlation functions (TCFs) (see Fig. 3). In constructing these functions, we compared the results from individual single-molecule data sets to check for consistency before averaging these together. Typically, ~35 individual single-molecule data sets were used to compute each function. In addition, we performed numerical simulations of the PDFs and TCFs using the master equation approach described in Sect. 3.4. Additional details of the numerical methods we employed are described in the SI.

**Figure 3.**
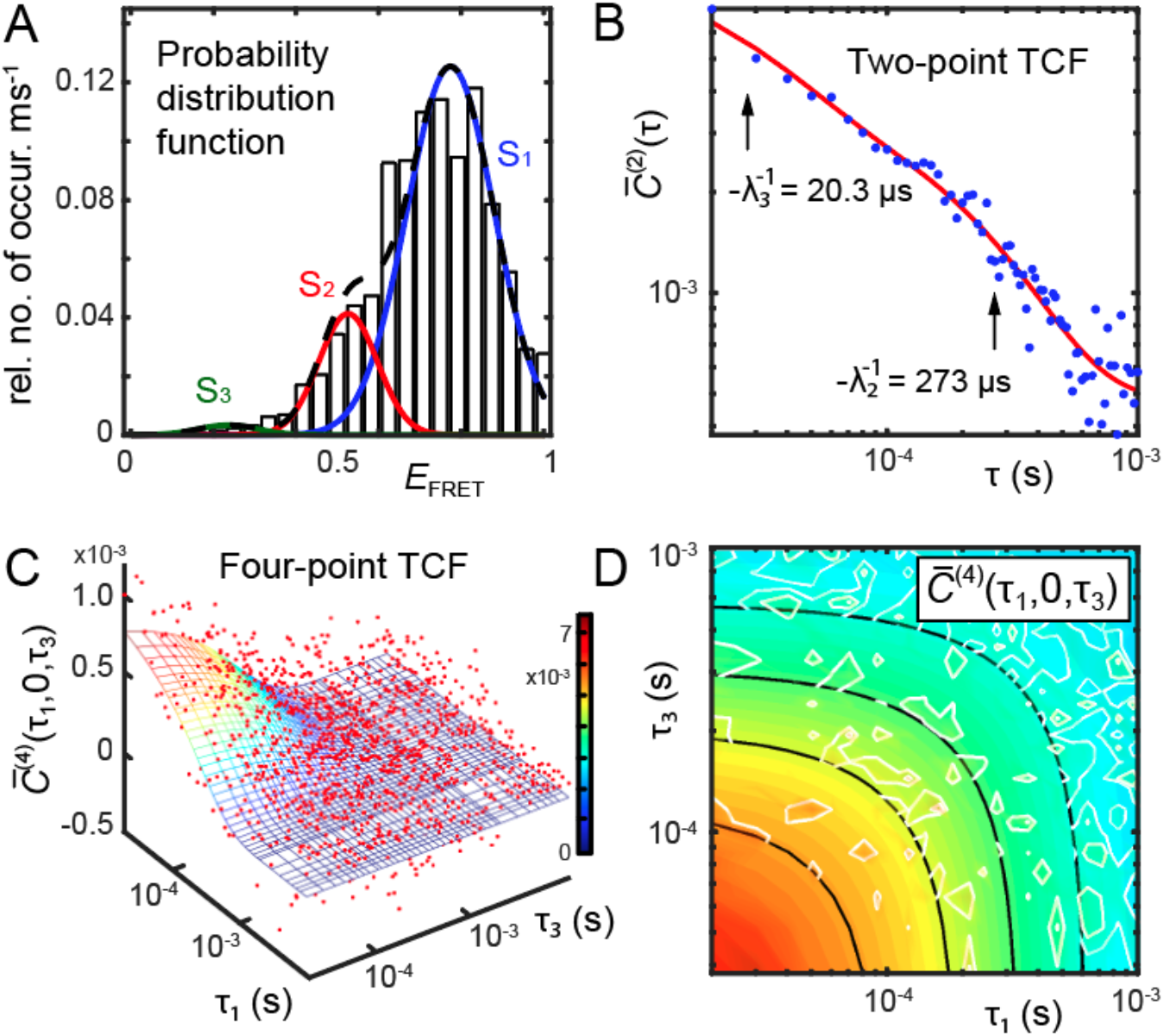
Experimentally derived statistical functions constructed from single-molecule photon correlation measurements of the 3’-Cy3/Cy5-oligo(dT)_15_-ss-dsDNA construct. Fits to model functions based on the kinetic master equation are also shown. (***A***) The normalized histogram of *E_FRET_* values using a sampling window *T_w_* = 1 ms and 21 million photons, which is represented as a sum of three Gaussian macrostates, labeled S_1_ – S_3_. (***B***) The two-point time correlation function (TCF) of *E_FRET_* values (blue points) is modeled theoretically as a weighted sum of two exponentially decaying terms (red curve) using a three-state master equation. (***C***) The four-point TCF is shown as a three-dimensional rendering (red points), which is overlayed with the theoretical model predicted by the three-state master equation (mesh surface). (***D***) The four-point TCF in comparison to the theoretical model are shown as twodimensional contour plots. The experimental data are shown as white contours and the model as a solid color-coded surface. The optimized values obtained from the master equation for the equilibrium PDF and the rate constant parameters are given in Table 2 and Table 3, respectively.

In Fig. 3*A*, we show a normalized histogram of *E_FRET_* values, which was compiled from the data streams of 35 individual single-molecule measurements of the 3’-Cy3/Cy5-oligo(dT)_15_-ss-dsDNA construct. The values of the FRET efficiency were calculated by binning the *D* and *A* intensities using a sampling window of *T_w_* = 1 ms. Each single-molecule photon data stream had an approximate mean flux *Ī_T_* ~20,000 s^−1^, and was collected over an acquisition period of 30 s. Thus, the total number of single-photon measurements used to compile the histogram is approximately ~21 million. The shape of the PDF, which is mostly peaked around *E_FRET_* ~ 0.7, but also exhibits a shoulder at *E_FRET_* ~ 0.5 and a tail for *E_FRET_* < 0.5, is consistent with our qualitative observations of the time-dependent behavior of the FRET efficiency trajectory shown in Fig. 2. The PDF cannot be modeled using a single Gaussian feature, but rather requires multiple underlying features to recover its shape. A spectral decomposition analysis suggests that the PDF is best modeled using a sum of three Gaussians, as indicated. All four of the ss-dsDNA constructs that we investigated exhibited a very similar shape to that shown in Fig. 3*A* (see Fig. S3 of the SI). We next discuss our kinetic analysis of these single-molecule data, which supports a model in which the system exists as a distribution of three quasi-stable macrostates in equilibrium. The results of this analysis provide optimized parameters for the Gaussian decomposition of the PDFs of all four of the ss-dsDNA constructs (Table S1 of the SI), and we list the results for the 3’-Cy3/Cy5-oligo(dT)_15_-ss-dsDNA construct in Table 2.

**Table 2.** Optimized parameters for the Gaussian decomposition of the FRET probabilit*y* distribution function (PDF) shown in Fig. 3*A*. Parameters defined in Eq. (2). Error bars for the mean FRET efficiencies are ~0.005 (see Fig. S11 and Table S10 of the SI).

| Construct | $A_1$ | $\langle E_{FRET} \rangle_1$ | $\sigma_1$ | $P_1^{eq}$ | $A_2$ | $\langle E_{FRET} \rangle_2$ | $\sigma_2$ | $P_2^{eq}$ | $A_3$ | $\langle E_{FRET} \rangle_3$ | $\sigma_3$ | $P_3^{eq}$ |
| --- | --- | --- | --- | --- | --- | --- | --- | --- | --- | --- | --- | --- |
| 3'oligo(dT)<sub>15</sub> | 3.04 | 0.77 | 0.11 | 0.81 | 1.00 | 0.52 | 0.07 | 0.18 | 0.08 | 0.23 | 0.07 | 0.01 |

We calculated experimental two-point TCFs [defined by Eq. (4)] from the time-dependent photon data streams of our single-molecule measurements (see SI for further details). For each of the ss-dsDNA constructs that we studied, we found that the two-point TCFs exhibited very similar decay dynamics, spanning the range of time scales: 10^-6^ – 10^-3^ s. These TCFs exhibited a relaxation component on the order of ~3 μs, as shown in Fig. S4 of the SI. We fit the first 20 microseconds of the TCFs to a model mono-exponential decay, and the optimized time constants for each of the ss-dsDNA constructs are listed in Table S2 of the SI. Relaxations on the time scale of ~3 μs have been observed previously in photophysical studies of the Cy3 chromophore, and are known to be associated with excited state intersystem crossing and forward photoisomerization, which undergo the reverse step of ground state recovery within ~10 μs (38–42). We therefore assign the ~ 3 μs decay in our current studies to photoisomerization of the Cy3 chromophore, and we assume that these relatively fast processes contribute to the ‘time-averaged’ background of the signal fluctuations that we observe on tens-of-microseconds and longer time scales, and which we interpret to be due to structural changes of the oligo(dT)_*n*_ templates.

In Fig. 3*B*, we show the experimental two-point TCF over the range of time scales from 2 ×10^-5^ – 10^-3^ s, which was calculated from the same 35 single-molecule measurements of the 3’-Cy3/Cy5-oligo(dT)_15_-ss-dsDNA construct used to compile the histogram shown in Fig. 3*A*. The decay is compared to a model bi-exponential fit, 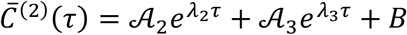, with relaxation time constants [eigenvalues of the rate matrix, Eq. (9)] 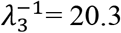 and 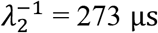, as shown. Upon modeling the experimental decay using a tri-exponential function, we obtained very similar values of the relaxation time constants, although the quality of the fit (as determined by the goodness-of-fit parameter *R*^2^) did not improve (see Fig. S5 and Table S3 of the SI). A comparison between the two-point TCFs and their model fits for all four ss-dsDNA constructs are shown in Fig. S6 of the SI, and the corresponding model fit parameters are listed in Table S4 of the SI.

We calculated the four-point TCFs from our experimental data according to Eq. (6). In Fig. 3*C*, we show a three-dimensional rendering of this function, 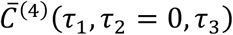, where we have set the intermediate time interval *τ*_2_ = 0. These data are shown overlayed with a two-dimensional model function, 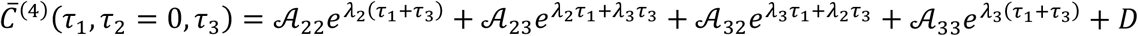, which is based on an optimized solution to the kinetic master equation. In Fig. 3*D*, we show the experimental and simulated four-point TCFs as a two-dimensional contour plot. We note that the three-dimensional surface of 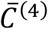 exhibits pronounced convex contour lines (corresponding to negative values of 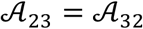, see Table S5), which indicate the presence of kinetic ‘bottleneck’ intermediates in the reaction pathway that inhibit the rapid exchange of population among the various macrostates of the system (19). We show the four-point TCFs and their model fits to all four ss-dsDNA constructs in Fig. S7 of the SI, and the corresponding model fit parameters are listed in Table S5 of the SI. The combined statistical functions shown in Fig. 3 serve to constrain a multi-parameter optimization of solutions to the master equation.

As discussed in Sect. 3.4, the presence of two decay components in the two-point TCFs implies that there must be at least three conformational macrostates between which the oligo(dT)_*n*_ templates interconvert. Moreover, a three-state model is consistent with the histogram of *E_FRET_* values (see Fig. 3*A*), in which the underlying Gaussian features, labeled S_1_ – S_3_, may be assigned to a ‘compact’ macrostate, a ‘partially-extended’ macrostate, and a ‘highly-extended’ macrostate, respectively. The compact macrostate S_1_ has the largest mean *E_FRET_* value, since it contains chain configurations with relatively short *D – A* chromophore distances. The partially-extended macrostate S2 and the highly-extended macrostate S3, on the other hand, have smaller mean *E_FRET_* values, indicating chain configurations with correspondingly larger *D – A* separations.

We therefore adopted a kinetic network scheme with *M* = 3 macrostates, as depicted in Fig. 4*A*, to model our results using the master equation [Eq. (8)]. The solutions to the three-state master equation can be written analytically in terms of the rate constants of elementary reactive steps (19). We performed numerical simulations of Eq. (8) by searching the parameter space of input rate constants and FRET efficiency values, as described in our prior work (14, 19). The solutions to the master equation are the time-dependent conditional probabilities *p_ij_*(*t*) [Eq. (12)] and the equilibrium distribution of states 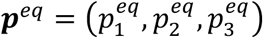 [Eq. (11)], which may be used to simulate expressions for the PDF and the two-point and four-point TCFs described in Sect. 3.3. We performed an iterative search of the parameter space in which the statistical functions *p*(*E_FRET_*), 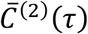 and 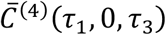, shown in Fig. 3 for the 3’-Cy3/Cy5-oligo(dT)_15_-ss-dsDNA construct, were calculated for a given set of input rate constants and FRET efficiency values, and compared to the experimentally-derived functions. The agreement between simulated and experimental functions was quantified using a linear least squares target function *χ*^2^, which was minimized using a multi-parameter regression algorithm to determine an optimized solution to the master equation (see SI for further details). We compared the results of our analyses using linear versus cyclical three-state network models, which showed that the cyclical model is favored. We confirmed that our optimized results represent a global minimum of the parameter space by monitoring the increase of the *χ*^2^ error function with variation of the time constant parameters (see Fig. S11 of the SI). The equilibrium populations and kinetic time constants resulting from of our analysis of the 3’-Cy3/Cy5-oligo(dT)_15_-ss-dsDNA construct are summarized in Table 2 and Table 3, respectively, and those of all four ss-dsDNA constructs are presented in Table S1 and Table S6 of the SI.

**Figure 4.**
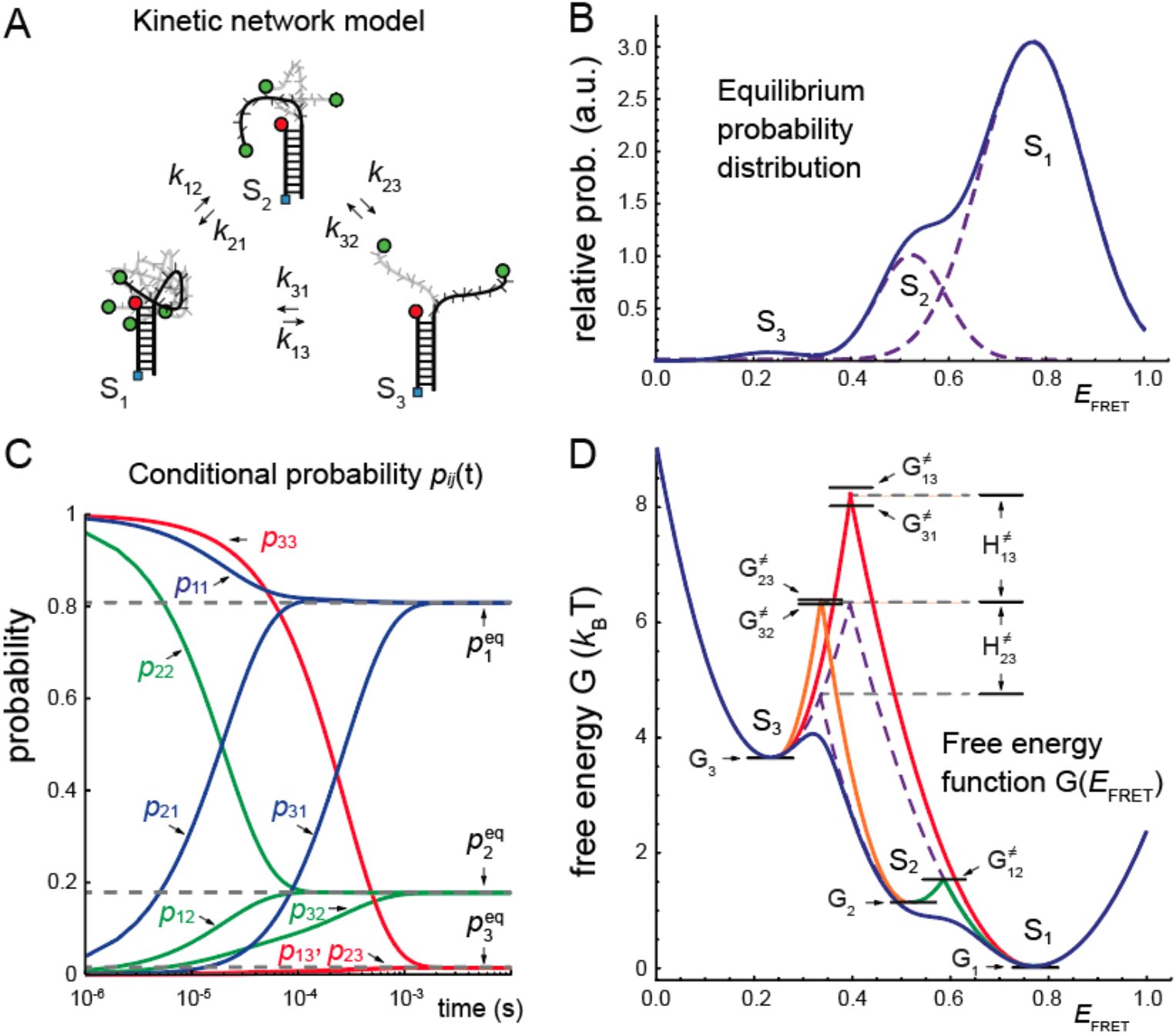
Theoretical modeling results of microsecond-resolved smFRET measurements of the 3’-Cy3/Cy5-oligo(dT)_15_-ss-dsDNA construct using the kinetic master equation. (***A***) Kinetic scheme depicting the elementary reactive steps connecting the three macrostates of the oligo(dT)_15_ template: ‘compact’ S_1_, ‘partially-extended’ S_2_, and ‘highly-extended’ S_3_. (***B***) The normalized probability distribution function (PDF) of *E_FRET_* values obtained from an optimized fit to the histogram shown in Fig. 3*A*. The distribution is represented as a sum of the *M* = 3 Gaussian macrostates with areas equal to the equilibrium probabilities 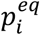, with *i* ∈ {1,2,3}. (***C***) The optimized solution to the master equation is the equilibrium PDF, ***p**^eq^*, and the time-dependent conditional probabilities, *p_ij_*(*t*), which give the likelihood that the system occupies the *j*th macrostate at time *t* if it previously occupied the *i*th macrostate at time 0, with *i,j* ∈ {1,2,3}. (***D***) The free energy landscape is constructed from the PDF shown in panel *B* according to the Boltzmann distribution [Eq. (2)]. Intersecting dashed purple lines indicate the entropic contributions to the free energy barriers that separate macrostates. Intersecting solid curves indicate the total free energy barriers calculated from the optimized rate constants according to the Arrhenius equation [Eq. (3)]. The optimized values obtained for the equilibrium PDF are given in Table 2, those for the rate constants are given in Table 3, and the free energy minima and transition barriers are given in Table 4. Estimates of the enthalpic and entropic contributions to the transition barriers are given in Table 5.

**Table 3.** Optimized forward and backward time constants corresponding to the elementary reactive steps obtained for the 3’-Cy3-Cy5-oligo(dT)_15_-ss-dsDNA construct. The time constant parameters are defined in Eq. (9). Error bars were determined from the cross-sections of the error function surfaces 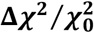, shown in Fig. S11 of the SI.

| Construct | $k_{12}^{-1}$ (ms) | $k_{21}^{-1}$(ms) | $k_{13}^{-1}$ (ms) | $k_{31}^{-1}$ (ms) | $k_{23}^{-1}$ (ms) | $k_{32}^{-1}$ (ms) |
| --- | --- | --- | --- | --- | --- | --- |
| 3'-oligo(dT)<sub>15</sub> | 0.113 ± 0.004 | 0.025 ± 0.0005 | 104 ± 1.85 | 1.85 ± 0.03 | 4.01 ± 0.03 | 0.325 ± 0.03 |

In Fig. 4*B*, we show the model equilibrium PDF obtained from our optimization analysis of the 3’-Cy3/Cy5-oligo(dT)_15_-ss-dsDNA construct. We have modeled the PDF as a sum of three component Gaussians, with normalized areas equal to the equilibrium populations of the three macrostates (see Table 2). Thus, macrostate S_1_ is the dominant component with 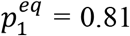 and mean FRET efficiency 〈*E_FRET_*〉_1_ = 0.77 ± 0.01, macrostate S_2_ is a minority component with 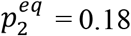 and mean FRET efficiency 〈*E_FRET_*〉_2_ = 0.52 ± 0.07, and macrostate S_3_ is a trace component with 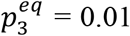 and mean FRET efficiency 〈*E_FRET_*〉_3_ = 0.23 ± 0.07. These optimized Gaussian parameters can be used to estimate the relative free energy minima of the macrostates using the Boltzmann relation [Eq. (2)].

In Fig. 4*D*, we plot the individual macrostate contributions to the free energy surface of the 3’-Cy3/Cy5-oligo(dT)_15_-ss-dsDNA construct, which are shown as dashed purple curves. In Fig. S8 of the SI, we compare the free energy surfaces for all four ss-dsDNA constructs. The sum of the macrostate contributions is shown as a solid blue curve. The three macrostates are represented by parabolas with minimum values 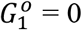 (taken as the ground state), 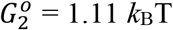 and 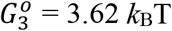. Here, the superscript ‘*o*’ indicates that we have taken the minimum of macrostate S_1_ as the reference for all other energies. The parabolic dependence of the free energy with respect to the *E_FRET_* parameter is due to our assumption of a Gaussian distribution of end-to-end distances of the oligo(dT)_*n*_ template for a given macrostate. The Gaussian distribution describes approximately the relative probability to observe a microscopic chain configuration with an end-to-end distance (and a corresponding *E_FRET_* value) that deviates from the mean value, and thus accounts for the entropic contribution to the free energy penalty associated with chain elongation or compression (10). The Gaussian model is, of course, accurate for long ideal polymer chains, and ignores the effects of excluded volume and other internal chain segment interactions. For the oligo(dT)_*n*_ template strands, the random walk statistics of chain configurations is additionally influenced by attractive and repulsive interactions between non-adjacent nucleotides and the tendency of adjacent bases to stack locally. Thus, the presence of three distinct macrostates, each with a different mean end-to-end distance, suggests that there exist chain configurations of the oligo(dT)_*n*_ templates with characteristically different local flexibilities, which is (presumably) due to the effects of different distributions of stacked bases. We note that in the context of the Gaussian macrostate model, the points at which the component parabolas (dashed purple curves) cross in the free energy surface approximate the entropic contributions to the transition state barriers for macrostate interconversion. However, the true free energy barriers are likely to include enthalpic contributions associated with the energy needed to rearrange the internal constraints imposed by a given distribution of stacked bases associated with a given macrostate. We next examine the free energy barriers of activation that result from the kinetic analysis of our data.

In Fig. 4*C*, we show the time-dependence of the optimized conditional probabilities *p_ij_*(*t*) where *i,j* ∈ {1,2,3}. These functions illustrate the kinetic behavior of the three-state system. The equilibrium populations are the asymptotic values approached by the conditional probabilities in the limit of long times. For example, the diagonal terms *p_ii_*(*t*) decay from unity to the equilibrium population value 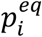 on the time scale 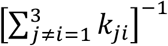, while off-diagonal terms *p_ji_*(*t*) increase from zero to the equilibrium population value 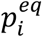 on the time scale 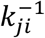. The relaxation dynamics of all nine conditional probability terms occur primarily on sub-millisecond time scales. The optimized values of the time constants for the 3’-Cy3/Cy5-oligo(dT)_15_-ss-dsDNA construct are presented in Table 3, and the time constants for all four of the ss-dsDNA constructs are listed in Table S6 of the SI. It is interesting to note that the fastest interconversion processes are between macrostates S_1_ and S_2_ (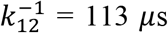 and 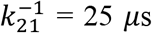), while the slowest interconversions are between macrostates S_1_ and S_3_ (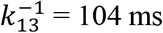 and 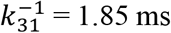) and between macrostates S_2_ and S_3_ (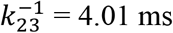 and 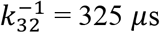).

From the optimized time constants, we determined the free energies of activation 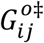 using the Arrhenius relation [Eq. (3)]. In calculating the various values for 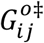, we have used the maximum rate constant resulting from our analysis 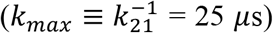 as a reference, which we assume corresponds to a barrierless downhill process for the transition S_2_ → S_1_ (i.e., 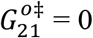). The remaining activation energies are measured relative to this reference. In Table 4, we list these activation energies for the 3’-Cy3/Cy5-oligo(dT)_15_-ss-dsDNA construct, and we compare the results of all four ss-dsDNA constructs in Table S7 of the SI. We plot these barrier heights as horizontal lines in the free energy diagram shown in Fig. 4*D*. It is interesting to note that the transition barrier corresponding to the uphill reaction S_1_ → S_2_ is 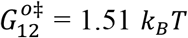 (green curve) is fully accounted for by the estimated entropic contribution 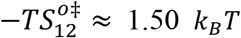 given by the intersection between the Gaussian components. We thus conclude that the activation barrier for the S_1_ → S_2_ transition is purely entropic such that 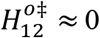. On the other hand, the transition barriers corresponding to the forward and backward reactions S_2_ ⇌ S_3_ are 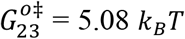 and 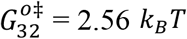, respectively. The absolute barrier heights corresponding to these values (5.08 + 1.11 = 6.19 *k_B_T* and 2.56 + 3.62 = 6.18 *k_B_T*, respectively), which are indicated on the free energy diagram in Fig. 4*D* (orange curve), exceed the estimated entropic contribution 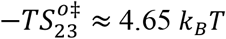, suggesting that the forward and backward transitions between macrostates S_2_ and S_3_ requires 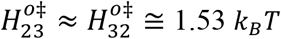 of internal redistribution energy. Similarly, the forward and backward transition barriers separating macrostates S_1_ and S_3_ are 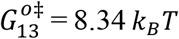 and 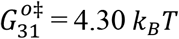, respectively. The absolute barrier heights corresponding to these values (8.34 + 0 = 8.34 *k_B_T* and 4.30 + 3.62 = 7.92 *k_B_T*, respectively) are plotted on the free energy surface shown in Fig. 4*D* as a red curve, and exceed the estimated entropic contribution 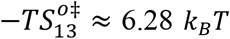, suggesting that the enthalpic contribution to the activation energies are 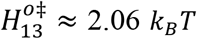 and 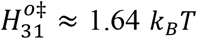. We list the estimated enthalpic and entropic contributions to the activation barriers of all four ss-dsDNA constructs in Table S8 of the SI. We note that these values are very similar for the four constructs, and the enthalpic contribution to the forward and backward activation barriers for both the S_1_ ⇌ S_3_ and the S_2_ ⇌ S_3_ processes are within the range ~1 - 3 *k_B_T*, which is of comparable magnitude to the enthalpy of base unstacking obtained for TT pairs (1.04 kcal mol^-i^) in thermal denaturation studies of nicked DNA (43).

**Table 4.** Optimized free energy minima and transition barriers for the 3’-Cy3/Cy5-oligo(dT)_15_-ss-dsDNA construct. The S_1_ macrostate is taken to be the ground state and excited state energies are given by 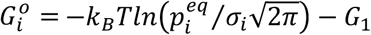 [Eq. (2)] with parameters 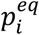 and *σ_i_* listed in Table 2. The activation energies are determined by 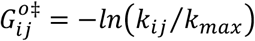 [Eq. (3) of the main text, with *k_max_* = *k*_21_ the fastest time constant of the system]. The rate constant parameters are listed in Table S6. Energies are listed in units of *k_B_T*.

| Construct | $G_1^o$ | $G_2^o$ | $G_3^o$ | $G_{12}^{o\ddagger}$ | $G_{21}^{o\ddagger}$ | $G_{13}^{o\ddagger}$ | $G_{31}^{o\ddagger}$ | $G_{23}^{o\ddagger}$ | $G_{32}^{o\ddagger}$ |
| --- | --- | --- | --- | --- | --- | --- | --- | --- | --- |
| 3'-oligo(dT)<sub>15</sub> | 0 | 1.11 | 3.62 | 1.52 | 0 | 8.34 | 4.30 | 5.08 | 2.56 |

## 5. Conclusions

In this paper we use microsecond-resolved smFRET experiments, recently developed in our laboratory, to begin to characterize conformational fluctuations in the structures of ssDNA at low concentrations in dilute aqueous solution. This is important for studies of protein-DNA interactions because – unlike dsDNA – the interactions stabilizing ssDNA conformations are not cooperative, and therefore conformations that exist at equilibrium, even at trace concentrations, are measurable with our present techniques. These results are likely to be significant from a biological perspective because our approach extends smFRET techniques to the microsecond time regime. By expanding the range of experimentally accessible time scales, it is thus possible to monitor not only the binding and unbinding of protein components of protein-DNA complexes on millisecond time scales and longer, but also to observe directly the interconverting conformational species (on sub-millisecond time scales) of ssDNA that comprise the interaction partners in the initial steps of biologically relevant protein-ssDNA interactions.

As described in the Introduction, and also in previous papers (14, 18, 19, 26), we have been using smFRET methods to study, in particular, the initial steps in the binding, assembly, and translocation of the T4 ssb proteins (gp32), and their cooperatively-bound clusters onto and along ssDNA lattices. Understanding the molecular details of these interactions with both the ssDNA lattices and ss-dsDNA junctions, which are present in the DNA constructs that we have studied, are central to obtaining molecular insight into the regulatory mechanisms of these ssb proteins as they interact with and control the functions of the other components of the bacteriophage T4 replisome – including the helicase and primosome, helicase loader proteins, the DNA polymerases and the processivity clamps and clamp loaders.

In the current paper, we describe microsecond-resolved studies on the ssDNA ‘tails’ of the ss-dsDNA constructs that we have used previously in our studies of the gp32 binding and assembly system (see Fig. 1). We studied short ss oligo-deoxythymidine [oligo(dT)_*n*_] templates of varying length (*n* = 14 and 15 nts) and polarity (3’ versus 5’, Table S1 of the SI) because gp32 binding and assembly onto these ss-dsDNA constructs depends sensitively on these factors. As we have shown, and as one might expect, these differences in the details of the constructs do not significantly change the equilibrium distribution of the template conformations, nor the dynamics of state-to-state interconversion. Nevertheless, it was necessary to investigate all four ss-dsDNA constructs to provide a baseline for the ssb interaction studies that will follow (37). Thus, the present study enables us to re-examine the ssb-ssDNA interaction systems with the ability to determine how ssb protein binding and assembly is directly coupled to the dynamics of the DNA interaction partners.

The oligo(dT)_*n*_ templates used in the present study comprised the ssDNA ‘tails’ of ss-dsDNA constructs, which had been labeled at their ends with a Cy3 donor chromophore and a Cy5 acceptor. We analyzed our data by constructing statistical functions; i.e., the equilibrium probability distribution function of FRET efficiency values and the two-point and four-point time correlation functions. We simulated these functions using a kinetic network model based on a master equation to obtain optimized parameters from which we have reached the following conclusions.

Our results indicate that the oligo(dT)_*n*_ templates of all four of the ss-dsDNA constructs used are dynamic systems with three distinct conformational macrostates: a ‘compact’ form S_1_, a ‘partially-extended’ form S_2_, and a ‘highly-extended’ form S_3_ (see Fig. 4*A*). These macrostates are separated in relative stability by just a few thermal energy units: 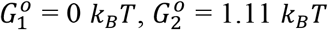, and 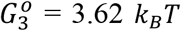 in the case of the 3’-Cy3-Cy5-oligo(dT)_15_-ss-dsDNA construct (see Fig. 4*D* and Table 4). The time scales of interconversion between these macrostates span four orders of magnitude, and range from tens-of-microseconds to hundreds-of-milliseconds (see Fig. 4*C* and Table 3). The fastest relaxation time that we observed 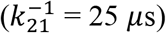 corresponds to the transition from the partially-extended macrostate S_2_ to the compact macrostate S_1_, which is commensurate with the relaxation time found for ssDNA loop closure in the formation of a DNA hairpin composed of an oligo(dT)_16_ loop (44, 45).

The relative stabilities and relatively long lifetimes of the three macrostates may be related to the energetic cost of unstacking and stacking adjacent bases. From the kinetic rate constants (Table 3) we determined the free energies of activation (Table 4), which we decomposed into entropic and enthalpic contributions (Table 5). The forward and backward enthalpies of activation that separate the highly-extended macrostate S_3_ from the partially-extended and compact macrostates, S_2_ and S_1_, respectively, have values within the range 1.5 to 2 *k_B_T*. If we consider the barriers for all four of the ss-dsDNA constructs (see Table S8 of the SI), the range of forward and backward enthalpy values increases from 1.2 to 3.1 *k_B_T*. We note that these values are comparable to the enthalpy of base unstacking for a single TT pair (1.04 kcal mol^-1^ = 1.73 *k_B_T*), which was obtained from thermal denaturation studies of the bent-stacked equilibrium in nicked DNA (43). Our estimates of the enthalpic contributions to the activation barriers suggest that only a small number of base stacking interactions are involved in these transformations.

**Table 5.** Transition barrier enthalpies and entropies for the 3’-Cy3/Cy5-oligo(dT)_15_-ss-dsDNA construct. The entropic contribution to the transition barrier is defined 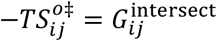 with 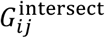 the free energy at the point of intersection between the parabolas describing macrostate-*i* and macrostate-*j* (dashed purple curves in Fig. 4*D*). The enthalpy of transition is estimated to be 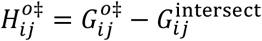. Enthalpic and entropic contributions to the transition barriers are given in units of *k_B_T*.

| Construct | $-TS_{12}^{o\ddagger}$ | $-TS_{13}^{o\ddagger}$ | $-TS_{23}^{o\ddagger}$ | $H_{12}^{o\ddagger}$ | $H_{21}^{o\ddagger}$ | $H_{13}^{o\ddagger}$ | $H_{31}^{o\ddagger}$ | $H_{23}^{o\ddagger}$ | $H_{32}^{o\ddagger}$ |
| --- | --- | --- | --- | --- | --- | --- | --- | --- | --- |
| 3'-oligo(dT)<sub>15</sub> | 1.50 | 6.28 | 4.65 | 0 | 0 | 2.06 | 1.64 | 1.54 | 1.53 |

The notion that multiple extended conformations of short oligomeric ssDNA templates exist as stable macrostates at equilibrium differs from our previous assumptions about short ssDNA lattices, in which only a single, relatively compact macrostate has been considered to be the primary conformation (14). The parameters that we have determined to characterize the free energy surface suggest a new model for ssDNA structure may be required (see Fig. 5). In order to gain additional insight into the nature of the extended macrostates, we carried out a coordinate transformation (24) of the free energy surface as a function of the FRET efficiency (shown in Fig. 4*D*) to one as a function of the donor – acceptor separation, which we equate with the end-to-end distance, *R_ee_*, of the oligo(dT)_15_ template (Fig. 5*A*). In performing this transformation, we used a value of the Förster transfer distance *R*_0_ = 56 Å obtained by others, which assumes that the relative orientation of the Cy3 donor and Cy5 acceptor, which are attached to the ss-dsDNA constructs using flexible linkers, is random and that the orientation factor *κ*^2^ = 2/3 (31, 32). These assumptions are further validated by our own recent smFRET experiments using polarized light excitation and detection (46). From the transformed free energy surface, we see that the local minima of the three macrostates, S_1_ – S_3_, may be assigned to mean end-to-end distances 〈*R_ee_*〉 = 46.5, 56.0 and 67.5 Å, respectively.

**Figure 5.**
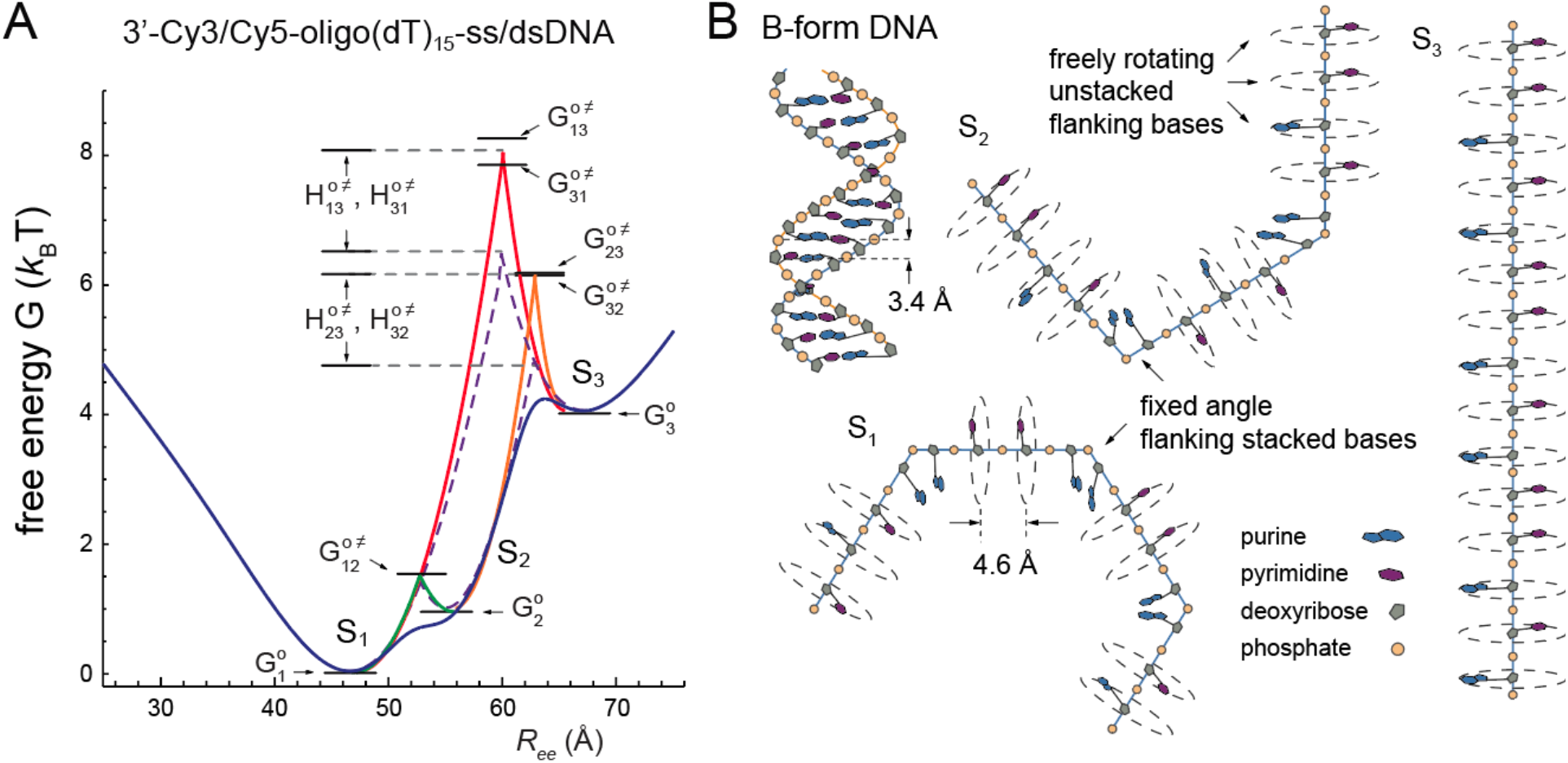
(***A***) Transformed free energy surface as a function of the end-to-end separation, *R_ee_*, of the oligo(dT)_15_ template. The transformation is given by *P*(*R_ee_*) = *P*(*E_FRET_*)*dE_FRET_*/*dR_ee_* with 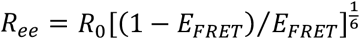 and *R*_0_ = 56 Å, and assumes that the relative orientation between the FRET Cy3 donor and Cy5 acceptor is random such that the orientation factor *κ*^2^ = 2/3 (31, 32). (***B***) Schematic microstate configurations of the oligo(dT)_15_ template exemplifying the three macrostates S_1_ – S_3_ described by the free energy surface shown in panel (***A***). In B-form DNA (upper left), the mean distance (rise per residue) between adjacent bases is 3.4 Å. In the oligo(dT)_15_ template strands (labeled S_1_ – S_3_), adjacent bases are primarily unstacked with a mean inter-base distance of ~4.6 Å. Unstacked bases are stabilized by the gain in entropy associated with the resultant enhanced rotation about the segmental axes. A sequence of unstacked bases forms a contiguous linear segment. The presence of a single stacked base ‘defect site,’ introduces a ‘vertex’ with an accompanying change in the bond vector.

We next consider how the extended macrostates S2 and S3 might play a biological role as intermediates for protein binding mechanisms (47, 48). For example, the T4 ssDNA binding (ssb) protein may be able to access its 7 nt binding site footprint from extended macrostate conformations, S_2_ and S_3_, more readily than from the compact macrostate S_1_. It is known from electron microscopy studies of cooperatively bound ssb nucleoprotein filaments that adjacent nucleotide bases within the complexes are fully unstacked with an average inter-base separation of 4.6 Å (5). This is in contrast to the 3.4 Å separation between stacked bases in B-form duplex DNA. The oligo(dT)_14_ and oligo(dT)_15_ templates examined in this study were chosen specifically to permit two ssb proteins to bind cooperatively within the final-formed nucleoprotein complexes. In the case of the 3’-Cy3/Cy5-oligo(dT)_15_-ss-dsDNA construct, the contour length of the ssDNA component within a cooperatively-bound (ssb)2-oligo(dT)_15_ filament is expected to be approximately ~15 × 4.6 Å = 69 Å. Given that the mean end-to-end distance corresponding to macrostate S3 (*R_ee_* = 67.5 Å) is only slightly smaller than the contour length of the fully unstacked oligo(dT)_15_ template strand, we suggest that the extended macrostates do indeed represent potential loading sites for ssb nucleoprotein filament assembly.

In light of our findings, we propose a new model for ssDNA secondary structure, which is depicted schematically in Fig. 5*B*. We posit that at physiological conditions, an isolated ssDNA template exists at thermal equilibrium with a significant fraction of its flanking bases unstacked. In the B-form DNA duplex, the highly ordered secondary structure of the double-helix is stabilized by cooperative short-range attractive interactions (stacking) between flanking bases (depicted as blue purines and red pyrimidines) and the network of Watson-Crick hydrogen bonds between opposing bases of the complementary single-strands. The duplex is marginally destabilized by the Coulomb repulsions between negatively charged phosphates (shown as yellow circles) of the sugar-phosphate backbones, which are partially screened by the condensation layer of positively charged counterions in the immediate aqueous surroundings.

We conjecture that adjacent stacked bases within an isolated segment of ssDNA, which lack the stabilizing influence of the complementary strand of duplex DNA, are only marginally stabilized with respect to thermal fluctuations that disrupt the short-range (solvophobic) attraction between base surfaces (49). Unstacked bases within an isolated segment of ssDNA are energetically favored by the Coulombic repulsion between adjacent phosphates and the gain in entropy associated with enhanced rotational freedom about the segmental axes. A sequence of metastable unstacked bases forms an essentially linear contiguous segment, which may be disrupted by the spontaneous formation of a stacking interaction between two adjacent bases along the ssDNA chain. The presence of such a vicinal stacked pair of bases introduces a ‘defect site,’ which then induces a change in the bond vector of two adjoining segments, each composed of unstacked bases. In Fig. 5*B*, we depict three examples of such microstate ssDNA conformations, which may each be assigned to one of the three macrostates S_1_ – S_3_, as shown. Each macrostate can be viewed as a stable family of microstates that contains a unique, dynamically equilibrated distribution of stacked base ‘defect sites.’ The principal difference between the underlying distributions of the three macrostates is the mean number of stacked base defect sites, whose presence effectively disrupts the linearity of the oligo(dT)_15_ template, and thereby reduces the end-to-end distance. The mean number of defect sites is maximized for the most stable macrostate S_1_ (depicted in the figure having three defect sites). The ground state S_1_ minimum represents the end-to-end distance (〈*R_ee_*〉 = 46.5) corresponding to balance between stabilizing and de-stabilizing (entropic and enthalpic) interactions, which arise in the presence of the dynamically interconverting stacked base defect sites. We further note that in the context of this model, conformational transformations between any two of the three macrostates, S_1_ – S_3_, correspond to the loss or gain of a single stacked base defect site, which is fully consistent with the values that we have determined for the relative stabilities and transition enthalpy barriers of the macrostates.

The above model for ssDNA secondary structure can provide useful insights for the interpretation of future microsecond-resolved smFRET studies of the T4 bacteriophage ssb nucleoprotein filament assembly mechanism using these same ss-dsDNA constructs. The initial steps of ssb protein-filament assembly involve the association of a ssb (gp32) monomer with an exposed ssDNA template binding site (7 nts), which may form on sub-millisecond time scales as a contiguous segment of unstacked flanking bases. Additional functional steps in which ssb proteins can assemble into cooperatively-bound clusters, slide along a ssDNA template strand and disassemble, will be mediated by the nucleation and dissolution of flanking stacked base defect sites, as described by our model. We anticipate that the model can be further tested by performing microsecond-resolved smFRET experiments on ss-dsDNA constructs with ssDNA sequences of purine or of mixed base sequences.

## Acknowledgements

A.H.M. acknowledges useful discussions with Prof. Marina Guenza. The authors are also grateful to our laboratory colleagues in the Marcus and von Hippel groups for many useful discussions. This work was supported by grants from the National Institutes of Health General Medical Sciences (GM-15792 to A.H.M. and P.H.v.H.). B.I. and C.S.A. were supported as predoctoral trainees by an NIH-NIGMS Institutional Research Service Award in Molecular Biology and Biophysics (GM-07759). P.H.v.H. is an American Cancer Society Research Professor of Chemistry.

## Supporting Information

### 1. Materials and Methods

#### Sample Preparation

We studied four single-stranded (ss) – double-stranded (ds) DNA constructs with a ss oligo(dT)_*n*_ overhanging ‘tail’ region, which varied in length (*n* = 14 and 15) and polarity (3’ and 5’, see Fig. 1 and Table 1 of the main text). DNA samples were purchased at nanomole quantities as dehydrated solids composed of individual single-strands (Integrated DNA Technologies, *IDT*, Coralville, IA, USA). The samples were rehydrated in an aqueous buffer composed of 100 mM NaCl, 6 mM MgCl_2_, and 10 mM Tris (pH 8.0) to reach a concentration of 100 μM. The ssDNA solutions were annealed by first diluting aliquots of the 100 μM Cy3-containing (‘template’) strand in buffer to reach 150 nM concentration, and the complementary 100 μM Biotin/Cy5-containing (‘primer’) strand to reach 100 nM concentration. The two solutions were combined to achieve a 2:3 (p:t) number ratio to ensure each primer strand has an attached template. The combined solution was heated to 94°C for 2 minutes and allowed to cool gradually to room temperature 22°C. The annealed ss-dsDNA constructs were stored at 4°C between experiments, or alternatively frozen at −4°C during extended periods. The annealed ss-dsDNA solution was diluted 1000-fold before it was introduced into the microscope sample chamber, as described below.

#### Microfluidic Sample Chamber

Sample chambers were constructed from microscope slides, which were chemically modified using the procedure described by Chandradoss *et al*. (see Fig. S1*A*) (1). Sample chambers were thoroughly cleaned with detergent (Alconox), water, acetone, KOH, acidic piranha-solutions (Sulfuric Acid + Hydrogen Peroxide), and methanol. The piranha solution served to clean the surface of the slide and to reduce the silicon-oxide to Si-OH. The reduced fused-silica quartz surface was treated with a solution of 3-aminopropyl-trimethoxy silane + acetic acid in methanol, and subsequently with poly(ethylene glycol)-succinimidyl valerate (PEG-SVA) (Laysan Bio, Inc.) with a molecular weight of 5000 Da in 100 mM sodiumbicarbonate solution to passivate the surface against interactions with proteins and to prevent nonspecific binding of DNA. A small fraction of the PEG that coated the microscope slides were labeled with biotin. Neutravidin (Thermo Fisher Scientific), a de-glycosylated avidin protein tetramer with strong binding affinity to biotin (2), was used as a linker to bind the biotin attached to the slide to the biotin-covalently bound to the ends of the ss-dsDNA constructs.

#### Oxygen Scavenging and Triplet State Quenching Reagents

An oxygen scavenging solution consisting of glucose oxidase, catalase and glucose were used to sequester free oxygen in the solution. The solution was introduced into the sample cell before smFRET data were acquired. A series of chemical reactions results in the net loss of oxygen from the buffer (3). In the absence of oxygen scavenging buffer, dissolved oxygen in the solution will irreversibly react with the excited-state fluorophores leading to photobleaching of the sample. In addition, the triplet state quencher Trolox (Sigma Aldrich) was used to prevent ‘photo-blinking,’ which is a process in which the chromophore probes reversibly interconvert between excited singlet and triplet states (4). Because the lifetimes of the excited triplet states are on the order of milliseconds, the resulting slow recovery of the singlet ground state leads to a dramatic decrease in the fluorescence signal rate.

#### Microsecond-Resolved Single-Molecule (sm) FRET

We performed microsecond-resolved smFRET experiments on the ss-dsDNA constructs using instrumentation and procedures described previously (5, 6). In Fig. S1*B*, we show a schematic of the instrumental setup. The microfluidic sample chamber was positioned using a computer-controlled translation stage (NPS-XY-100A with NPS3330 controller, Queensgate, UK), which was mounted to an inverted fluorescence microscope (Nikon 2000 TE Eclipse). The sample was illuminated using a continuous wave 532 nm laser (Coherent Compass) in a prism-based total internal conversion fluorescence (TIRF) geometry. The incident laser was focused to a 50 μm diameter spot on the sample using a 100 mm plano-convex lens, and the ensuing fluorescence was collected using a 100 × oil-immersion objective (Nikon CF160, Plan Apochromat, N.A. 1.4, w.d. 130 μm). The fluorescence from a single molecule was transmitted through a 100 μm pinhole (ThorLabs), which was positioned at the downstream image plane of the 200 mm microscope lens tube. The transmitted signal was optimized by scanning the XY-stage with sub-nanometer precision. The fluorescence was collimated using a 75 mm biconvex lens, which was placed one focal length downstream from the pinhole. The fluorescence was directed toward a 635 nm dichroic beam splitter (FF640-FDi01-25×36, Semrock, USA), and the separated donor Cy3 and acceptor Cy5 fluorescence channels were focused onto avalanche photodiodes (APDs, SPCM AQR-13, Perkin-Elmer Optoelectronics, Singapore) using long working distance objective lenses (SLCPlanFl, 40x, N.A. 0.55, w.d. 2.6 mm, Olympus, Hauppauge, NY). To ensure that the molecules we studied were localized within a uniform and uncrowded region of the sample chamber, we used an electron multiplying charge-coupled device camera (emCCD, iXon DV897-BB, 512 × 512 pixels, Andor Technologies, Belfast, Northern Ireland) to acquire wide-field images at 30 ms resolution.

#### Data Acquisition

The laser power at the sample was adjusted to 10 mW, which resulted typically in a combined donor-acceptor fluorescence photon count rate of ~ 15,000 – 20,000 cps. The signal ‘background rate’ of ~ 2,000 cps was determined by scanning the XY-stage position away from the center of a molecule. Single-photon detection resulted in the emission of TTL voltage pulses from the APDs, which were sent to a custom built 64 MHz digital counter. The donor Cy3 and acceptor Cy5 signals were each composed of a stream of temporally separated TTL pulses, which were ‘time-stamped’ at a resolution of 0.1 μs using a 10 MHz digital counter and stored on a computer hard disk. A typical single-molecule data run was recorded for a thirty-second duration. In post-data acquisition, we determined the time-dependent *D*(*A*) signal rates according to *I*_*D*(*A*)_(*t*) = *N*_*D*(*A*)_(*t*)/*T_w_*, where *N*_*D*(*A*)_ was the number of *D*(*A*) photons detected during an adjustable dwell period *T_w_* (≥ 10 *μ*sec) at time *t*. Multiple single-molecule data sets were recorded, each for a thirty-second duration. These signal trajectories were initially screened to reject cases that contained photobleaching events or multiple *D* chromophores. From the time-dependent signal rates we determined the FRET efficiency according to: *E_FRET_*(*t*) = *I_A_*(*t*)/[*I_D_* (*t*) + *I_A_*(*t*)]. An example trajectory for the 3’-Cy3/Cy5-oligo(dT)_15_-ss-dsDNA construct is shown in Fig. S2 for an integration window of *T_w_* = 1 ms.

**Figure S1.**
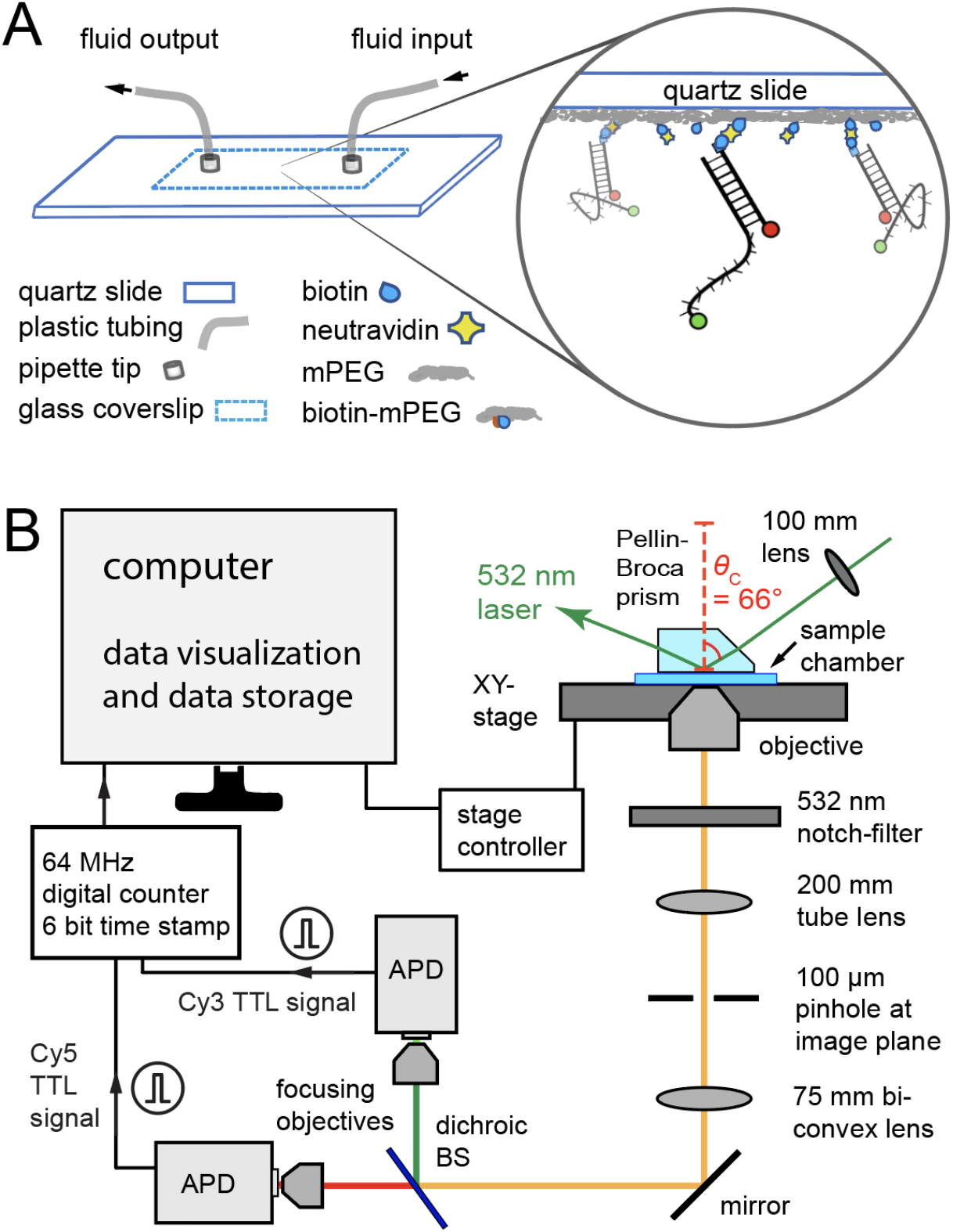
Experimental instrumentation used for microsecond-resolved single-molecule Förster resonance energy transfer (smFRET) experiments described the text. (***A***) Microfluidic sample chambers were constructed from fused-silica quartz microscope slides as described above. (***B***) The experimental setup is a modified version of the one described by Phelps *et al*. (5). APD = avalanche photodiode, BS = beam splitter, TTL = transistor-transistor logic, *θ_c_* = critical angle.

**Figure S2.**
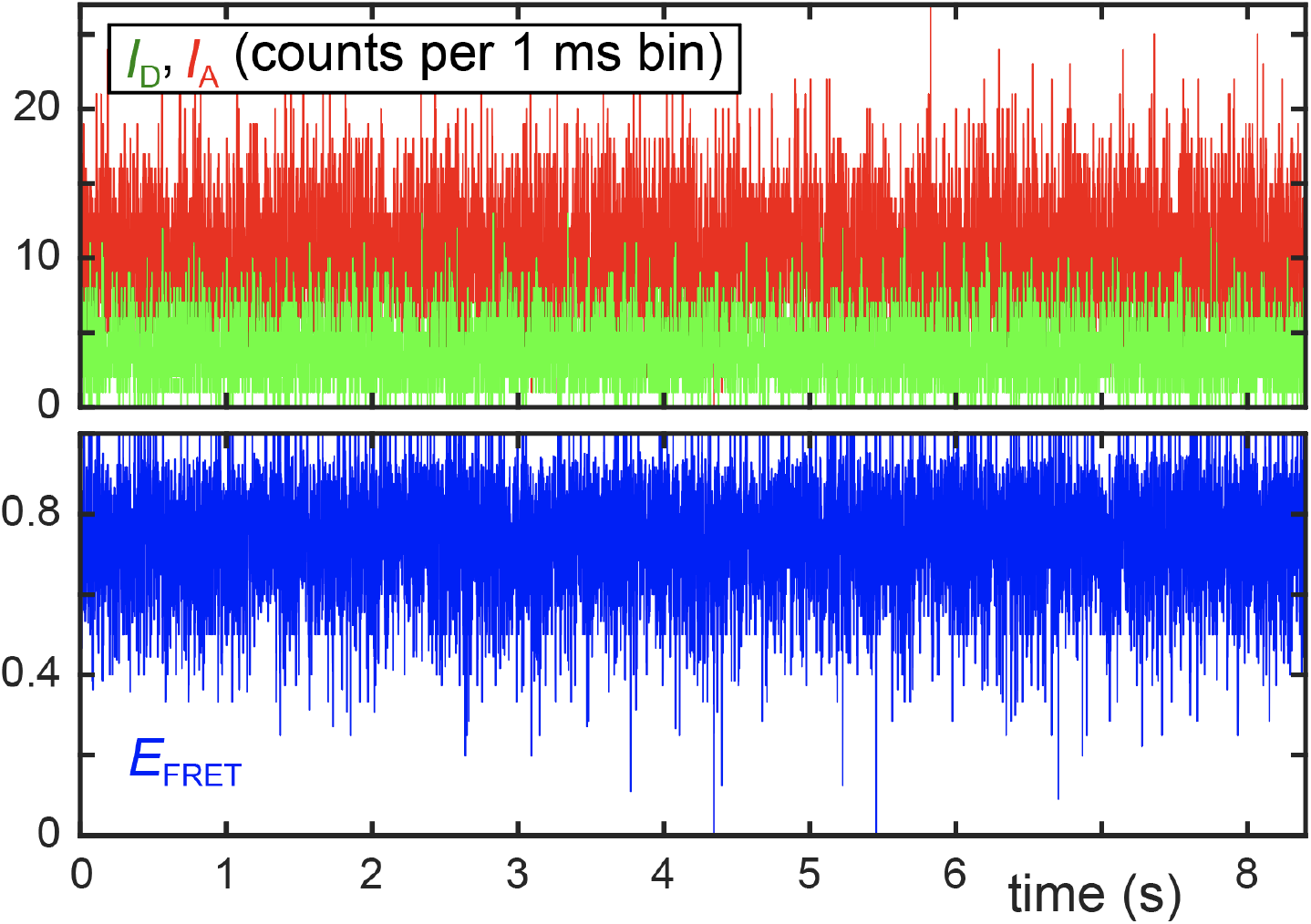
Example smFRET trajectory for the 3’-Cy3/Cy5-oligo(dT)_15_-ss-dsDNA construct. The donor *D* (Cy3, green) and acceptor *A* (Cy5, red) fluorescence intensities, *I*_*D*(*A*)_ (*t*) = *N*_*D*(*A*)_(*t*)/*T_w_*, are shown in the top panel with bin resolution *T_w_* = 1 ms. The FRET efficiency *E_FRET_*(*t*) = *I_A_*(*t*)/[*I_D_*(*t*) + *I_A_*(*t*)], which is calculated from the *I*_*D*(*A*)_ signals, is shown in the bottom panel (blue). The full data set was recorded for a duration of 30 seconds, although the trajectory is plotted here over a period of 8.5 seconds.

**Figure S3.**
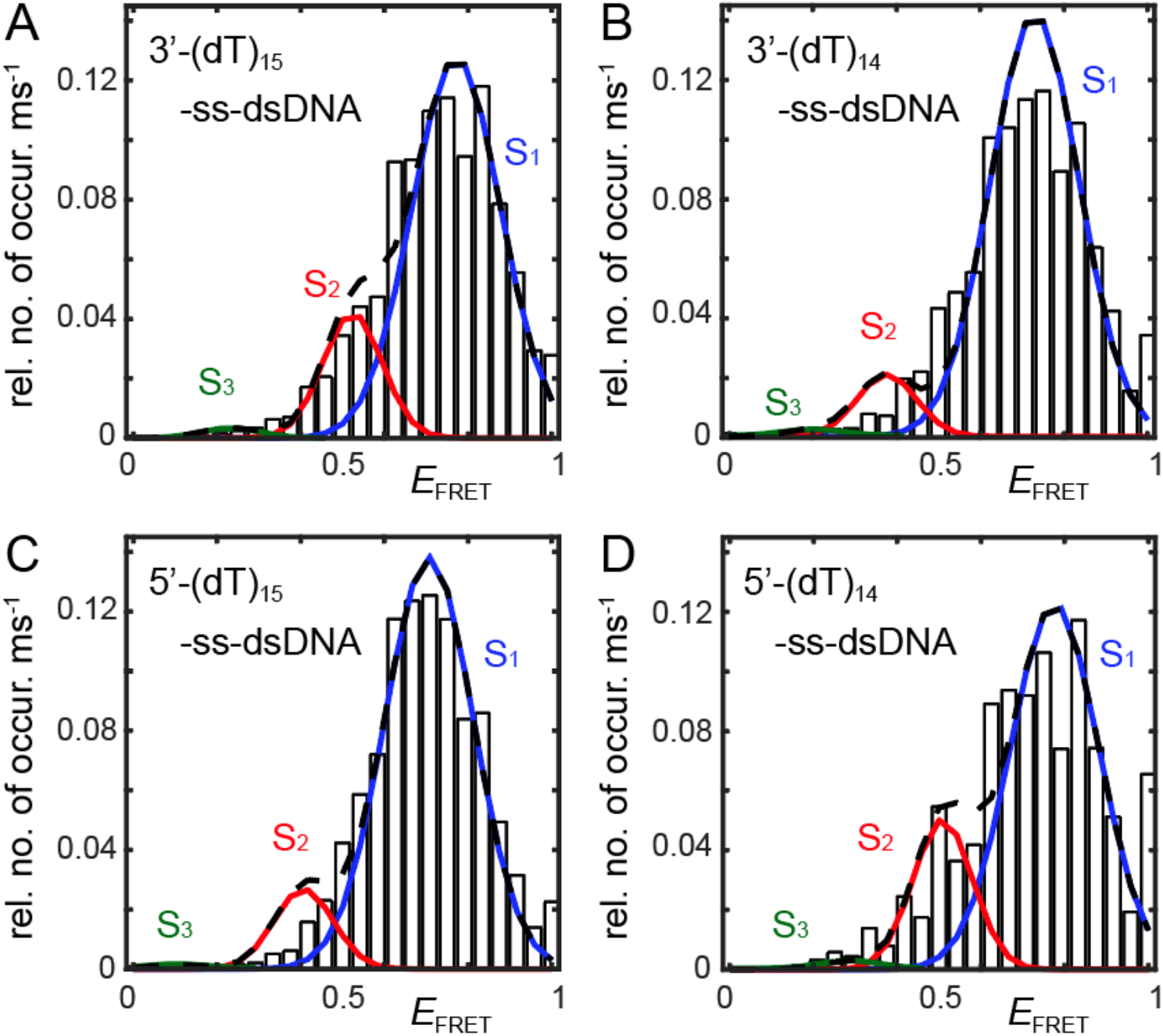
Experimental histograms and PDFs for the four ss-dsDNA constructs. The blue, red, and green Gaussian curves denote the compact (S_1_), partially extended (S_2_), and highly extended (S_3_) macrostate conformations, respectively. The optimized parameters obtained from our analysis of each of these data sets using the three-state master equation are listed in Table S1.

**Table S1.**
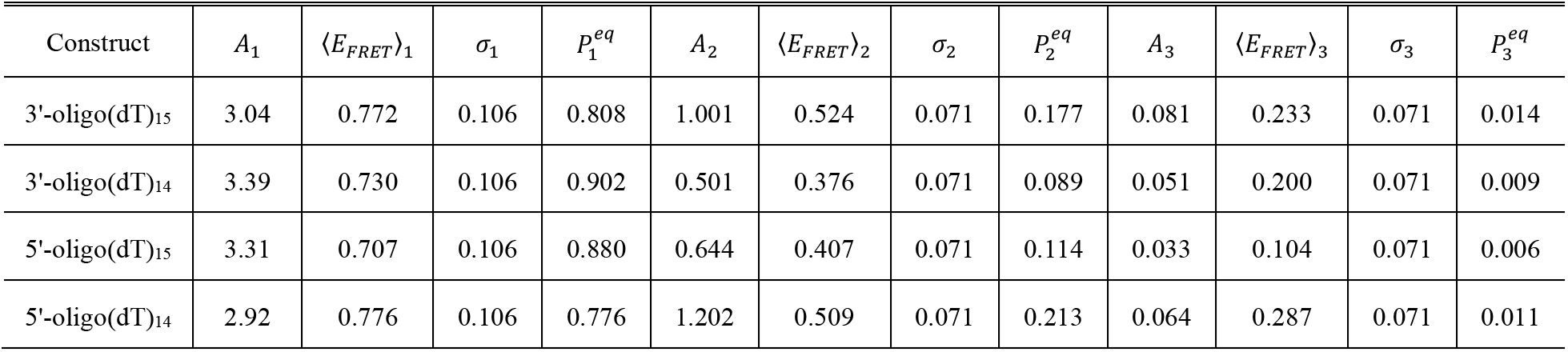
Optimized parameters corresponding to the FRET probabilit*y* distribution functions (PDFs) obtained for the four ss-dsDNA constructs studied in this work. The parameters are defined in the main text in and around Eq. (2), with 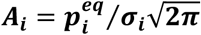. Error bars for the mean FRET efficiency values are on the order of the last significant figure, and are listed for the 3**’**-oligo(dT)_15_ construct in Table S11.

#### Numerical Calculations of Two-Point and Four-Point Time Correlation Functions (TCFs)

We calculated the two-point and four-point time correlation function (TCFs) for each single molecule data set. A typical microsecond-resolved single-molecule photon data stream contained ~30 million time points with a mean signal flux ~20,000 s^-1^. We compared the TCFs constructed from individual molecules for consistency before averaging them together to obtain the final result.

If the photon data stream is binned over a sufficiently long window *T_w_* (typically ~1 ms), the signal intensities, *I*_*D*(*A*)_(*t*) = *N*_*D*(*A*)_(*t*)/*T_w_* (where *N*_*D*(*A*)_ is the number of *D*(*A*) photons detected during the period *T_w_*) are vary continuously with time. In this case, the Wiener-Khinchin theorem can be applied to compute the two-point TCF according to:

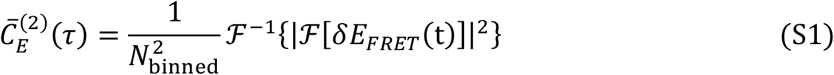

where 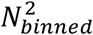 is the number of binned time points over which the FRET efficiency signal *δE_FRET_*(*t*) = *E_FRET_*(*t*) – 〈*E_FRET_*〉, with *E_FRET_*(*t*) = *I_A_*(*t*)/[*I_D_*(*t*) + *_A_*(*t*)], is monitored and 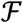 and 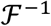 represent, respectively, the forward and inverse fast-Fourier-transform (FFT).

Because our microsecond-resolved smFRET signal is sparsely sampled at a much higher frequency than the mean signal flux, the data stream is non-continuous and we cannot use Eq. (S1) to calculate the two-point TCFs. Instead, we implemented the following pair-wise point-by-point multiplication scheme:

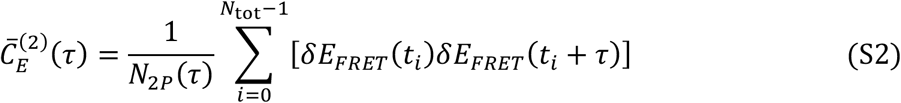

In Eq. (S2), the two-point TCF is a rolling time-average of the product of two successive measurements, *δE_FRET_*(*t_i_*)*δE_FRET_*(*t_i_* + *τ*), which are spaced apart by the interval *τ*. Here, *t_i_* is the measurement time index, *N*_tot_ (≈ 30 million) is the total number of evenly spaced data points, and *N*_2*P*_(*τ*) is the number of non-zero terms that contribute to the sum. An important consequence of the sparse data sampling of these measurements is that the number *N*_2*P*_(*τ*) depends non-uniformly on the interval τ. By accounting for the precise number of non-zero terms that contribute to the sum, it is possible to determine accurately the amplitude and shape of the two-point TCF. We have compared the results of Eq. (S2) to those using the Wiener-Khinchin formula [Eq. (S1)] for cases in which the signal varies continuously in time. We note that in this limit the two methods provide equivalent results, as expected.

To determine the four-point TCFs, we implemented a point-by-point multiplication scheme similar to the one we used for the two-point TCFs:

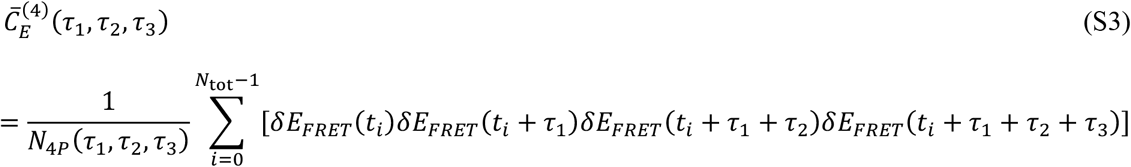

In Eq. (S3), *N*_4*P*_(*τ*_1_, *τ*_2_, *τ*_3_) is the number of non-zero four-point product terms that contribute to the sum, which are spaced apart by the intervals *τ*_1_, *τ*_2_ and *τ*_3_. For all calculations of the four-point TCFs presented in this work, we have set the intermediate time interval *τ*_2_ = 0.

The experimental two-point and four-point TCFs do not fully decay to zero at long times, but rather decay to non-zero asymptotic values. To account for this, we included an empirically-determined constant offset in all of our decay models. For the two-point TCFs, we used the model 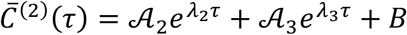, and for the four-point TCFs, we used the model 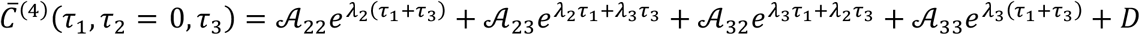. The values that we used for the constant offsets of the two-point and four-point TCFs (*B* and *D*, respectively) were determined from the means of the last 3 calculated data points of each surface, corresponding to times at which the decays had reached their asymptotic values.

#### Instrument Response Function

We performed control measurements on a concentrated solution of Rhodamine 6G (Rh6G), which fluoresces over a broad spectral range that spans both the donor and acceptor emission channels. Successive photon emission events from the concentrated Rh6G solution were uncorrelated on the 0.1 μs resolution of the instrument. The TCF of these data were used to construct the instrument response function (IRF), which has full-width-at-half-maximum ~ 1 μs. We used the IRF in a deconvolution procedure (7) to accurately determine the fastest time scale components (microseconds) of the TCFs that we determined from our smFRET measurements of the ss-dsDNA constructs (see Fig. S4 and Table S2).

**Figure S4.**
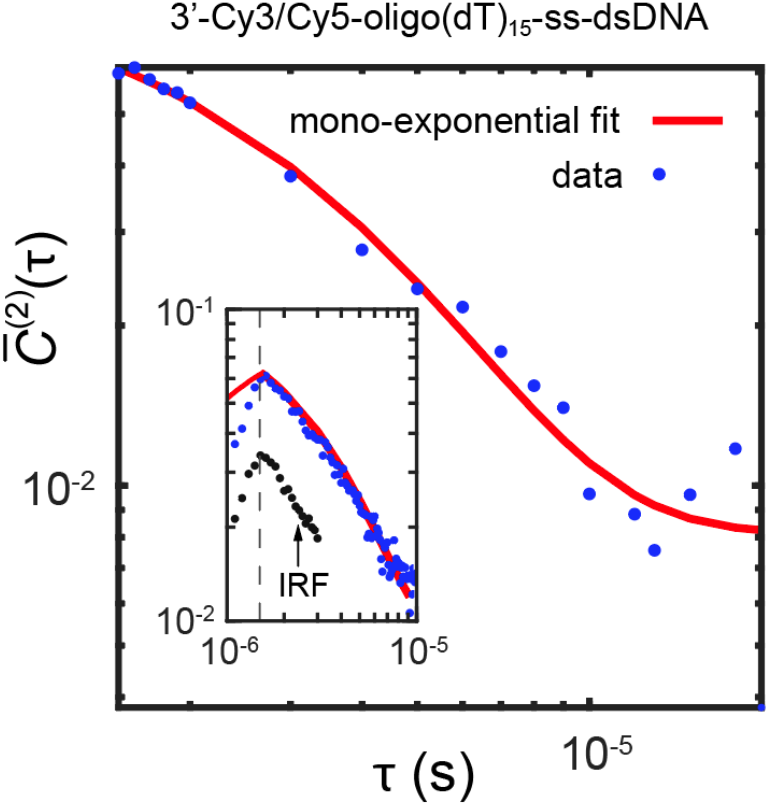
Two-point TCF of the 3’-Cy3/Cy5-oligo(dT)_15_-ss-dsDNA construct is shown over the range of time scales: 1.5 × 10^-6^ - 2 × 10^-5^ s. The first 20 μs of the autocorrelation function was fit to a model monoexponential decay using the equation *C*^(2)^ = *Ae*^−*t*/*τ*_0_^ + *B*, where *B* is a constant y-axis offset. The inset shows the measured instrument response function used to perform the short-time fits. All four of the ss-dsDNA constructs exhibited a single exponential decay constant ~3 μs (see Table S2), which we interpret to be an effect of the Cy3 excited state photoisomerization process.

**Table S2.** Decay constants determined from mono-exponential fits to the two-point TCFs over the range of time scales: 10^-6^ - 2 × 10^-5^ s. Each of the four ss-dsDNA constructs exhibited a decay with time constant ***τ*_0_** ≈ 3 μs, as shown for the case of the 3’-Cy3/Cy5-oligo(dT)_15_-ss-dsDNA construct in Fig. S4.

| Construct | Time Constant $\tau_0$ ($\mu\text{s}$) |
| --- | --- |
| 3'-oligo(dT)<sub>15</sub> | $2.9 \pm 0.45$ |
| 3'-oligo(dT)<sub>14</sub> | $2.7 \pm 0.35$ |
| 5'-oligo(dT)<sub>15</sub> | $3.2 \pm 0.51$ |
| 5'-oligo(dT)<sub>14</sub> | $3.0 \pm 0.50$ |

**Figure S5.**
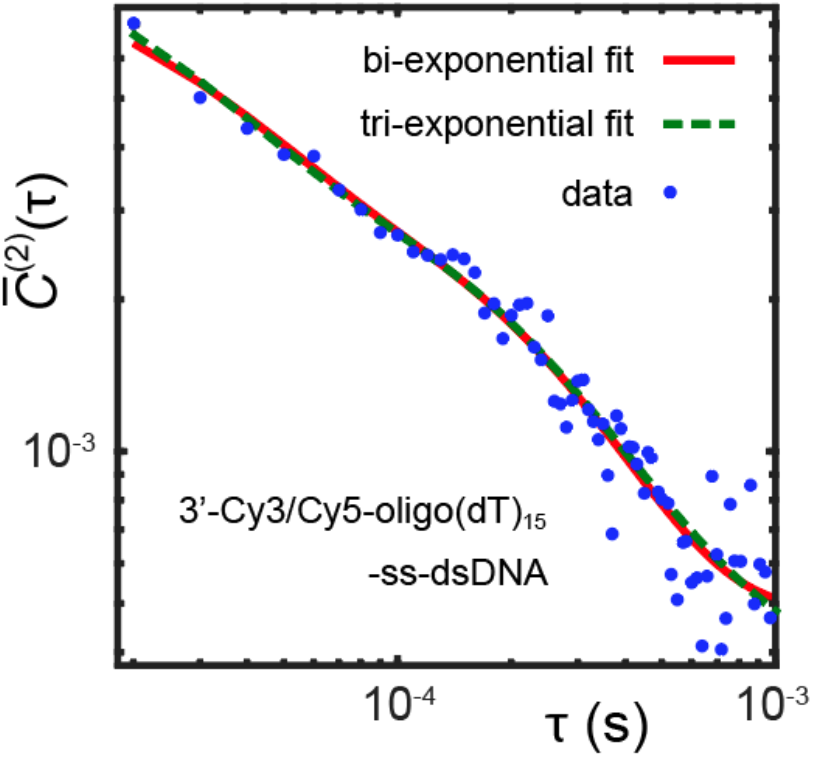
Comparison between model fits of the experimental two-point TCF to bi-exponential and triexponential functions. The bi-exponential function describes the decay to an acceptable level of accuracy.

**Table S3.** Results of numerical regression analysis of fitting multi-exponential model functions to experimental two-point TCFs for the four different ss-dsDNA constructs investigated in this work. The model functions are sums of exponentials: 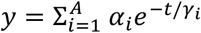, where *A* = 1, 2 or 3 to find the *α_i_* and *γ_i_* that maximize the *R*^2^ value. An example comparison between the best fits of a bi-exponential (*A* = 2) and triexponential (*A* = 3) model function to the experimental two-point TCF of the 3’-Cy3/Cy5-oligo(dT)_15_-ss-dsDNA construct is shown in Fig. S5. The values listed are the goodness-of-fit parameter *R*^2^. For each of the ss-dsDNA constructs, the *R*^2^ value does not improve significantly upon increasing the complexity of the model function from a bi-exponential to a tri-exponential decay.

| Construct | Mono-exponential | Bi-exponential | Tri-exponential |
| --- | --- | --- | --- |
| 3'-oligo(dT)<sub>15</sub> | 0.939 | 0.980 | 0.982 |
| 3'-oligo(dT)<sub>14</sub> | 0.912 | 0.963 | 0.968 |
| 5'-oligo(dT)<sub>15</sub> | 0.925 | 0.937 | 0.944 |
| 5'-oligo(dT)<sub>14</sub> | 0.900 | 0.934 | 0.937 |

**Figure S6.**
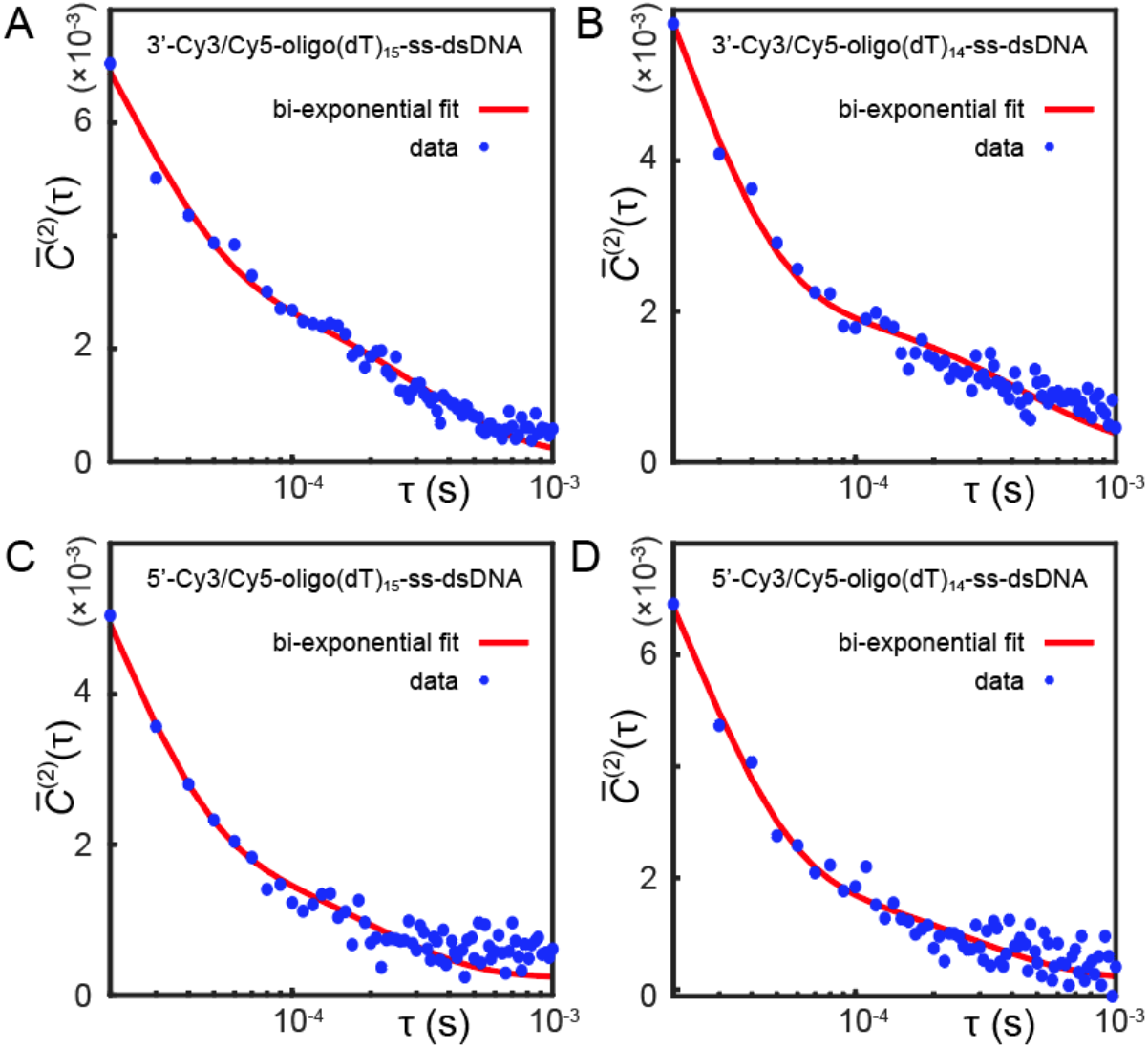
Two-point TCFs of the four ss-dsDNA constructs. (***A***) 3’-Cy3/Cy5-oligo(dT)_15_; (***B***) 3’-Cy3/Cy5-oligo(dT)_14_; (Q 5’-Cy3/Cy5-oligo(dT)_15_; and (***D***) 5’-Cy3/Cy5-oligo(dT)_14_. Red curves are model biexponential functions 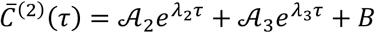, which are based on the optimized values obtained using the three-state master equation (see Table S4).

**Table S4.**
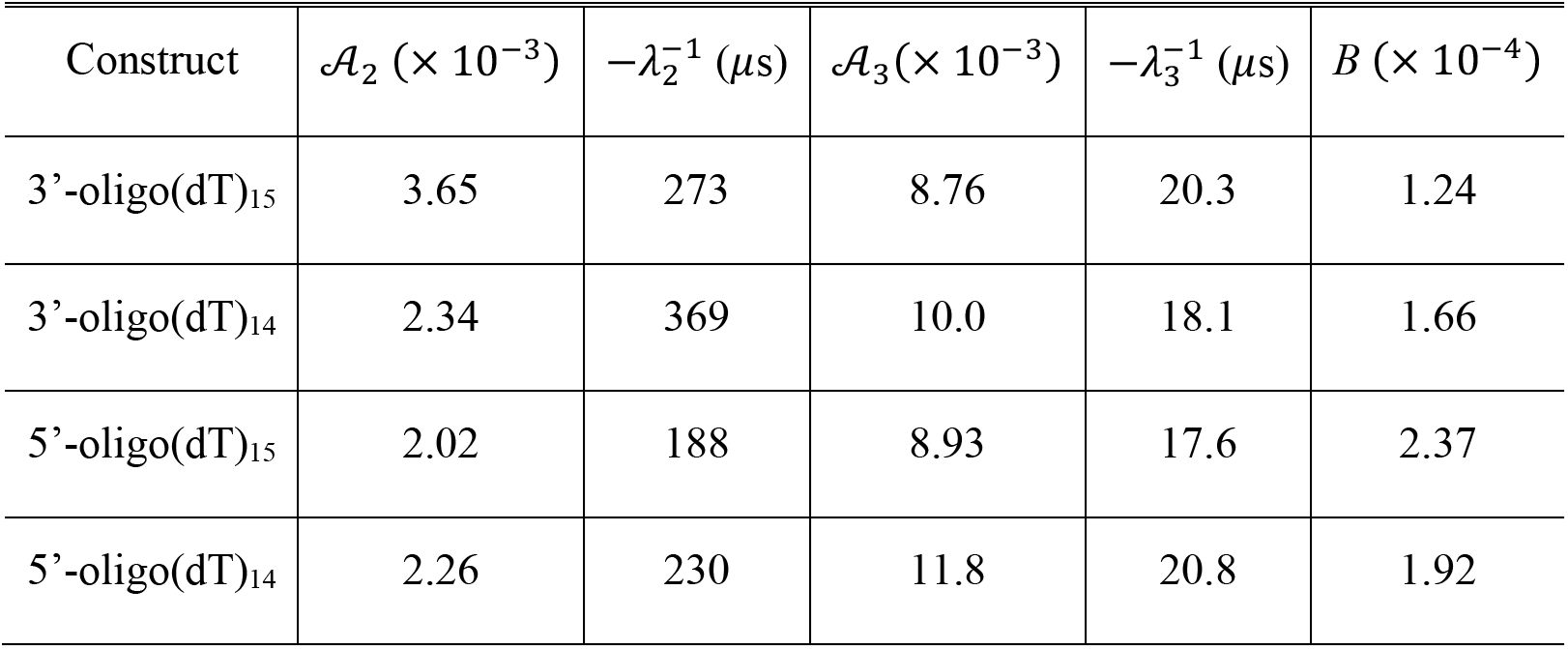
Optimized parameters for the model bi-exponential functions 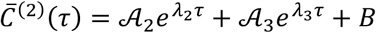 corresponding to the red curves shown in Fig. S6.

| Construct | $\mathcal{A}_2 (\times 10^{-3})$ | $-\lambda_2^{-1} (\mu s)$ | $\mathcal{A}_3 (\times 10^{-3})$ | $-\lambda_3^{-1} (\mu s)$ | $B (\times 10^{-4})$ |
| --- | --- | --- | --- | --- | --- |
| 3'-oligo(dT)<sub>15</sub> | 3.65 | 273 | 8.76 | 20.3 | 1.24 |
| 3'-oligo(dT)<sub>14</sub> | 2.34 | 369 | 10.0 | 18.1 | 1.66 |
| 5'-oligo(dT)<sub>15</sub> | 2.02 | 188 | 8.93 | 17.6 | 2.37 |
| 5'-oligo(dT)<sub>14</sub> | 2.26 | 230 | 11.8 | 20.8 | 1.92 |

**Figure S7.**
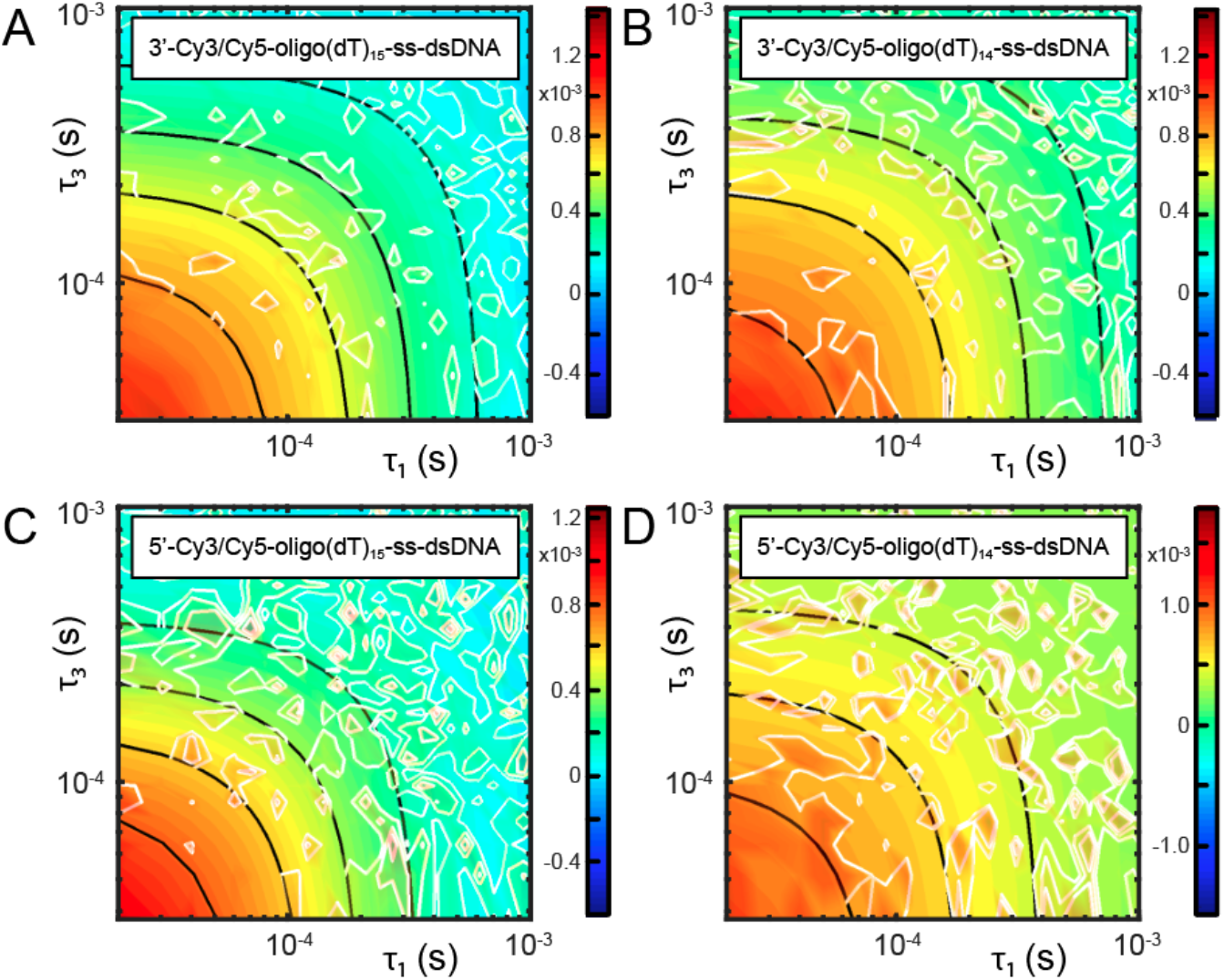
Overlay of experimental and simulated four-point TCFs of the four ss-dsDNA constructs, which are shown as two-dimensional contour plots. (***A***) 3’-Cy3/Cy5-oligo(dT)_15_; (***B***) 3’-Cy3/Cy5-oligo(dT)_14_; (***C***) 5’-Cy3/Cy5-oligo(dT)_15_; and (***D***) 5’-Cy3/Cy5-oligo(dT)_14_. Black curves are model functions 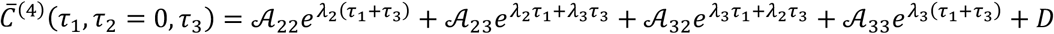, which are based on the optimized values obtained using the three-state master equation (see Table S5).

**Table S5.** Optimized parameters for the model functions 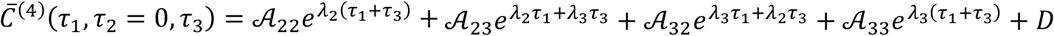, corresponding to the black contour lines shown in Fig. S7. Note that the eigenvalues, *λ*_2_ and *λ*_3_, are the same as those listed in Table S4.

| Construct | $\mathcal{A}_{22} (\times 10^{-4})$ | $\mathcal{A}_{33} (\times 10^{-4})$ | $\mathcal{A}_{23} = \mathcal{A}_{32} (\times 10^{-5})$ | $D (\times 10^{-5})$ |
| --- | --- | --- | --- | --- |
| 3'-oligo(dT)<sub>15</sub> | 8.53 | 2.82 | -2.86 | 5.56 |
| 3'-oligo(dT)<sub>14</sub> | 5.65 | 9.21 | -2.15 | 5.30 |
| 5'-oligo(dT)<sub>15</sub> | 6.40 | 5.46 | -2.79 | 1.26 |
| 5'-oligo(dT)<sub>14</sub> | 4.07 | 3.98 | -2.17 | 1.35 |

**Table S6.** Optimized forward and backward time constants corresponding to the elementary chemical steps obtained for the four ss-dsDNA constructs studied in this work. The time constant parameters are defined in Eq. (9) of the main text. Error bars are on the order of the last significant figure, and are listed for the 3’-oligo(dT)_15_ construct in Table S11.

| Construct | $k_{12}^{-1}$ (ms) | $k_{21}^{-1}$ (ms) | $k_{13}^{-1}$ (ms) | $k_{31}^{-1}$ (ms) | $k_{23}^{-1}$ (ms) | $k_{32}^{-1}$ (ms) |
| --- | --- | --- | --- | --- | --- | --- |
| 3'-oligo(dT)<sub>15</sub> | 0.113 | 0.025 | 104 | 1.85 | 4.01 | 0.325 |
| 3'-oligo(dT)<sub>14</sub> | 0.203 | 0.020 | 155 | 1.55 | 4.80 | 0.488 |
| 5'-oligo(dT)<sub>15</sub> | 0.154 | 0.020 | 273 | 1.80 | 4.15 | 0.211 |
| 5'-oligo(dT)<sub>14</sub> | 0.097 | 0.027 | 137 | 2.00 | 4.92 | 0.262 |

**Figure S8.**
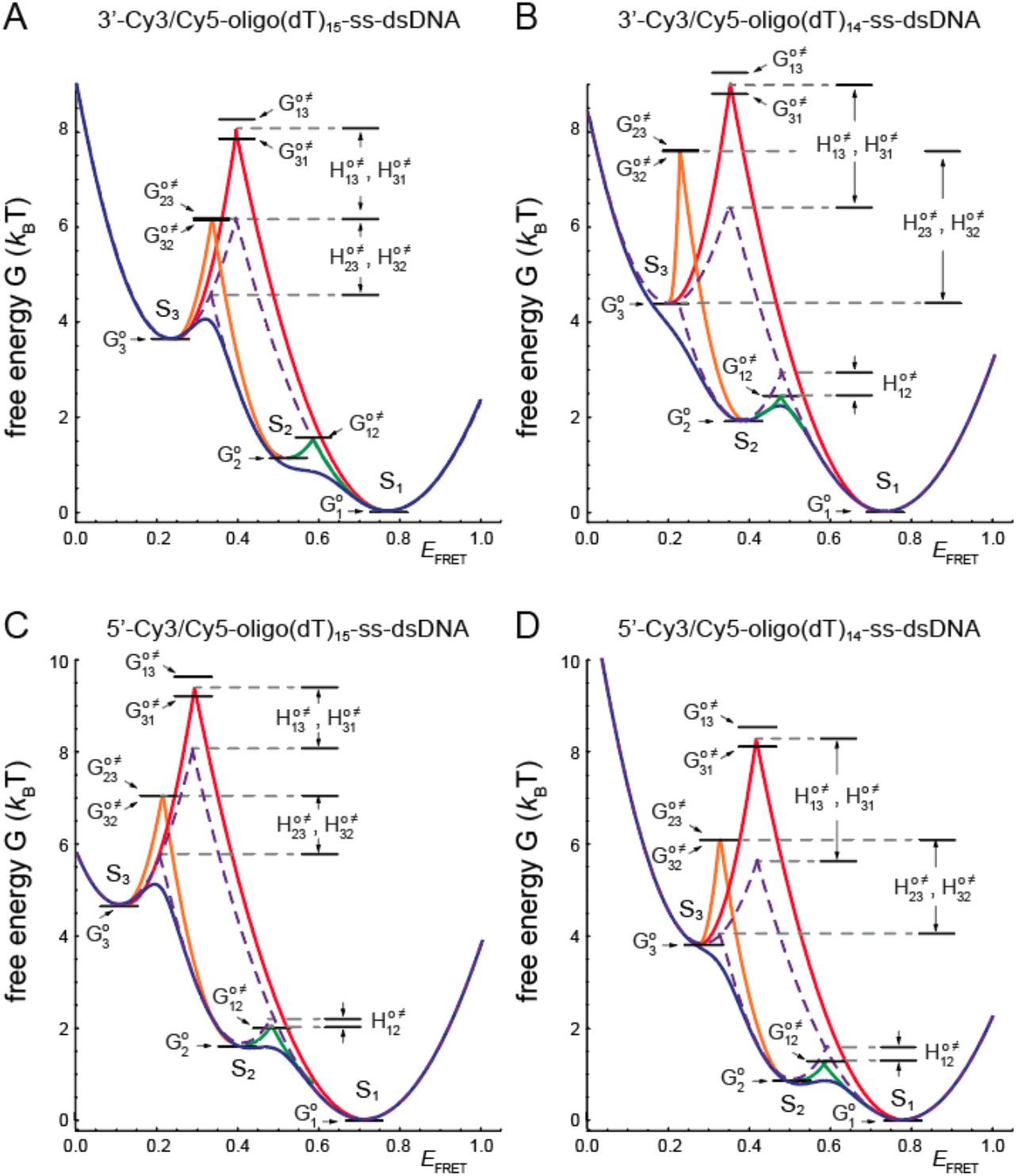
Free energy landscapes constructed from microsecond-resolved smFRET experiments for the four ss-dsDNA constructs: (***A***) 3’-Cy3/Cy5-oligo(dT)_15_; (***B***) 3’-Cy3/Cy5-oligo(dT)_14_; (***C***) 5’-Cy3/Cy5-oligo(dT)_15_; and (***D***) 5’-Cy3/Cy5-oligo(dT)_14_. The free energy landscape is constructed from the PDFs (Fig. S3 and Table S1) according to the Boltzmann distribution [Eq. (2) of the main text]. Intersecting dashed purple lines indicate the entropic contributions to the free energy barriers that separate macrostates. Intersecting solid curves indicate the total free energy barriers calculated from the optimized rate constants (Table S6) according to the Arrhenius equation [Eq. (3) of the main text]. Values of the free energy minima and transition barriers are given in Table S7. Estimates to the enthalpic and entropic contributions to the transition barriers are given in Table S8.

**Table S7.** Optimized free energy minima and transition barriers for the four ss-dsDNA constructs studied in this work. The S_1_ macrostate is taken to be the ground state and excited state energies are given by 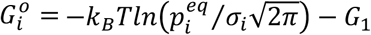 [Eq. (2) of the main text] with parameters 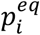 and *σ_i_* listed in Table S1. The activation energies are determined by 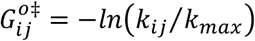 [Eq. (3) of the main text, with *k_max_* = *k*_21_ the fastest time constant of the system]. The rate constant parameters are listed in Table S6. Energies are listed in units of *k_B_T*.

| Construct | $G_1^o$ | $G_2^o$ | $G_3^o$ | $G_{12}^{o\ddagger}$ | $G_{21}^{o\ddagger}$ | $G_{13}^{o\ddagger}$ | $G_{31}^{o\ddagger}$ | $G_{23}^{o\ddagger}$ | $G_{32}^{o\ddagger}$ |
| --- | --- | --- | --- | --- | --- | --- | --- | --- | --- |
| 3'-oligo(dT)<sub>15</sub> | 0 | 1.11 | 3.62 | 1.52 | 0 | 8.34 | 4.30 | 5.08 | 2.56 |
| 3'-oligo(dT)<sub>14</sub> | 0 | 1.91 | 4.20 | 2.32 | 0 | 8.96 | 4.35 | 5.48 | 3.19 |
| 5'-oligo(dT)<sub>15</sub> | 0 | 1.64 | 4.62 | 2.04 | 0 | 9.52 | 4.50 | 5.34 | 2.36 |
| 5'-oligo(dT)<sub>14</sub> | 0 | 0.89 | 3.82 | 1.28 | 0 | 8.53 | 4.30 | 5.20 | 2.27 |

**Table S8.** Transition barrier enthalpies and entropies for interconversion between macrostates S_1_, S_2_ and S_3_. Entropies The entropic contribution to the transition barrier is defined 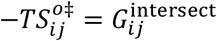 with 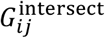 the free energy at the point of intersection between the parabolas describing macrostate-*i* and macrostate-*j* (dashed purple curves in Fig. 4*D* of the main text). The enthalpy of transition is estimated to be 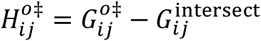. Enthalpic and entropic contributions to the transition barriers are given in units of *k_B_T*.

| Construct | $-TS_{12}^{o\ddagger}$ | $-TS_{13}^{o\ddagger}$ | $-TS_{23}^{o\ddagger}$ | $H_{12}^{o\ddagger}$ | $H_{21}^{o\ddagger}$ | $H_{13}^{o\ddagger}$ | $H_{31}^{o\ddagger}$ | $H_{23}^{o\ddagger}$ | $H_{32}^{o\ddagger}$ |
| --- | --- | --- | --- | --- | --- | --- | --- | --- | --- |
| 3'-oligo(dT)<sub>15</sub> | 1.50 | 6.28 | 4.65 | 0.02 | 0 | 2.06 | 1.64 | 1.54 | 1.53 |
| 3'-oligo(dT)<sub>14</sub> | 2.90 | 6.45 | 4.25 | -0.58 | 0 | 2.51 | 2.10 | 3.14 | 3.14 |
| 5'-oligo(dT)<sub>15</sub> | 2.25 | 7.95 | 5.70 | -0.21 | 0 | 1.57 | 1.17 | 1.28 | 1.28 |
| 5'-oligo(dT)<sub>14</sub> | 1.55 | 5.65 | 4.00 | -0.27 | 0 | 2.88 | 2.47 | 2.09 | 2.09 |

**Figure S9.**
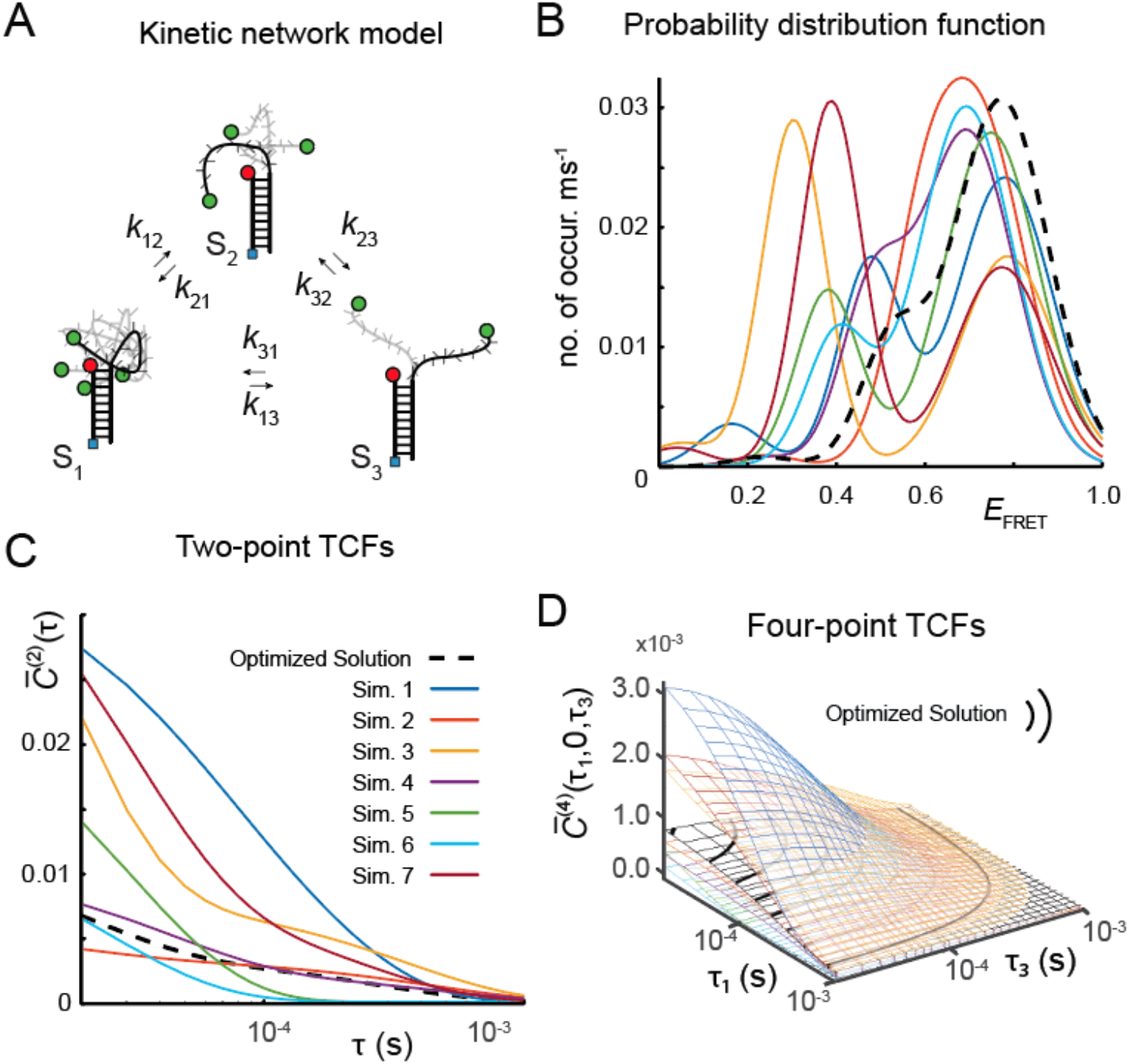
Comparison between the data-optimized solution to the three-state master equation for the 3’-Cy3/Cy5-oligo(dT)_15_-ss-dsDNA construct and various non-optimized solutions (labeled simulations). (***A***) Kinetic scheme depicting the elementary chemical steps connecting the three macrostates of the oligo(dT)_15_ template: ‘compact’ S_1_, ‘partially-extended’ S_2_, and ‘highly-extended’ S_3_. (***B***) Simulated probability distribution functions (PDFs). (***C***) Simulated two-point time correlation functions (TCFs). (***D***) Simulated four-point TCFs. The comparison serves to demonstrate the sensitivity of the analysis to the choice of model input parameters. The values of the time constant parameters and macrostate FRET efficiencies used for the simulations are listed in Table S9.

**Table S9.** Parameters used for the simulated PDFs and TCFs shown in Fig. S9.

| Trial | $k_{12}^{-1}$ (ms) | $k_{21}^{-1}$ (ms) | $k_{13}^{-1}$ (ms) | $k_{31}^{-1}$ (ms) | $k_{23}^{-1}$ (ms) | $k_{32}^{-1}$ (ms) | $\langle E_{FRET} \rangle_1$ | $\langle E_{FRET} \rangle_2$ | $\langle E_{FRET} \rangle_3$ |
| --- | --- | --- | --- | --- | --- | --- | --- | --- | --- |
| Opt. Sol. | 0.113 | 0.025 | 104 | 1.85 | 4.01 | 0.325 | 0.772 | 0.524 | 0.233 |
| Sim. 1 | 0.148 | 0.070 | 12.5 | 1.24 | 0.742 | 0.155 | 0.780 | 0.476 | 0.164 |
| Sim. 2 | 0.086 | 0.021 | 160 | 3.41 | 5.21 | 0.452 | 0.716 | 0.585 | 0.238 |
| Sim. 3 | 0.028 | 0.031 | 32.8 | 2.43 | 5.51 | 0.372 | 0.786 | 0.303 | 0.047 |
| Sim. 4 | 0.173 | 0.056 | 175 | 3.40 | 7.57 | 0.457 | 0.695 | 0.478 | 0.238 |
| Sim. 5 | 0.118 | 0.042 | 129 | 2.35 | 5.86 | 0.030 | 0.748 | 0.380 | 0.108 |
| Sim. 6 | 0.125 | 0.031 | 155 | 0.05 | 4.07 | 0.051 | 0.693 | 0.402 | 0.140 |
| Sim. 7 | 0.054 | 0.067 | 54.8 | 3.47 | 4.70 | 0.244 | 0.774 | 0.388 | 0.040 |

#### Numerical Calculations using the Three-State Master Equation

We adopted a kinetic network model that is described by the master equation [Eq. (8) of the main text] with the number of macrostates *M* = 3. We assumed a cyclical kinetic scheme, as shown in Fig. S9*A*. The six forward and backward rate constants obey the detailed balance condition (8, 9)

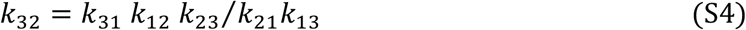

In order to model the statistical functions constructed from our data (two-point and four-point TCFs and PDFs), it is necessary to determine the six rate constants in addition to the values of each of the macrostate’s mean FRET efficiencies, 〈*E_FRET_*〉_*i*_ with *i* ∈ {1,2,3}, which leads to a total of eight independently adjustable parameters in our model.

The rate matrix, ***K*** [Eq. (9) of the main text] is written:

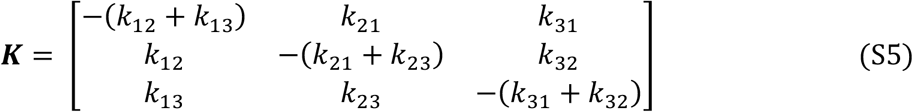

The eigenvalues and eigenvectors of ***K*** provide the relaxation times and collective modes of the coupled system of elementary chemical reactions. The eigenvalues and eigenvectors of ***K*** are used in Eq. (12) to determine the state-to-state conditional probabilities, *p_ij_*(*t*) with *i,j* ∈ {1,2,3}, and the equilibrium probability, 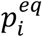, for each macrostate. The equilibrium probabilities, together with the mean FRET efficiencies, were used to determine the PDF according to: 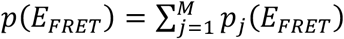 [see discussion around Eq. (2)]. The conditional probabilities and the mean FRET efficiencies were used in Eq. (5) and Eq. (7) to calculate the two-point and four-point TCFs, respectively. In Fig. S9*B*, we show a set of simulated PDFs for randomly chosen sets of rate constants and mean FRET efficiencies (listed in Table S9). Simulated two-point and four-point TCFs corresponding to the same sets of parameters are shown in Fig. S9*C* and Fig. S9*D*, respectively. The comparison between the simulated functions and those corresponding to the optimized solutions (shown as solid black dashed curves in Fig. S9) serves to demonstrate the sensitivity of this analysis to the choice of model input parameters.

#### Solving the Master Equation using the Similarity Transform Method

We applied a computationally efficient means to obtain solutions to the master equation by using the similarity transformation discussed in Sec. 3.4 (10):

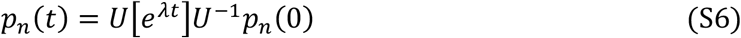

In Eq. (S6), the unitary matrix ***U*** = [***v***_1_, ***v***_2_,…, ***v**_M_*] has columns that are the eigenvectors of the rate matrix ***K***. The matrix [***e**^**λ**t^*] is diagonal with non-zero elements [*e^λt^*]_*ii*_ = *e^λ_i_t^*, where the decay constants *λ*_1_, *λ*_2_ and *λ*_3_ are the eigenvalues of ***K***.

After imposing the condition *λ*_1_ = 0 (discussed in Sec. 3.4 of the main text), the matrix [***e^λt^***] can be written:

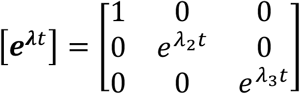

We rewrite Eq. (S6) in terms of the conditional probability matrix ***p***(*t*) = [***p***_1_, ***p***_2_,…, ***p**_M_*]:

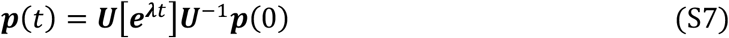

Upon setting the initial condition matrix ***p***(0) equal to the identity matrix ***I***, which represents all possible initial conditions, the matrix elements of ***p***(*t*) become the conditional probabilities *p_ij_*(*t*) according to:

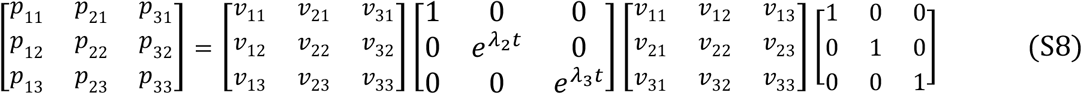

#### Iterative Multidimensional Optimization Procedure

We performed an iterative multi-parameter optimization procedure, similar to the one described by Phelps *et al*. (6, 9), to find the set of kinetic and thermodynamic parameters that best matches the experimentally-derived PDFs and two-point and four-point TCFs.

We introduce as a metric the weighted error function, *χ*^2^, to assess the quality of agreement between the master equation solutions and the statistical functions we have constructed from our experimental data. We define the error function as the cumulative squared-difference between the measured and simulated quantities, represented as *y* and *g*, respectively:

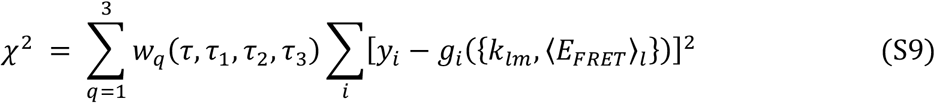

In Eq. (S9), *q* is an index that enumerates: (1) the FRET histogram; (2) the two-point TCF; and (3) the four-point TCF, and *i* is the domain index for each function. We chose the weighting coefficients, *w_n_*, so that the histogram and the TCFs contributed approximately equally to the error function. In addition, *w_n_* is a function of the lag times *τ, τ*_1_, *τ*_2_, *τ*_3_, such that the TCFs are assigned more weight at earlier times, as we did in our previous studies (6).

**Figure S10.**
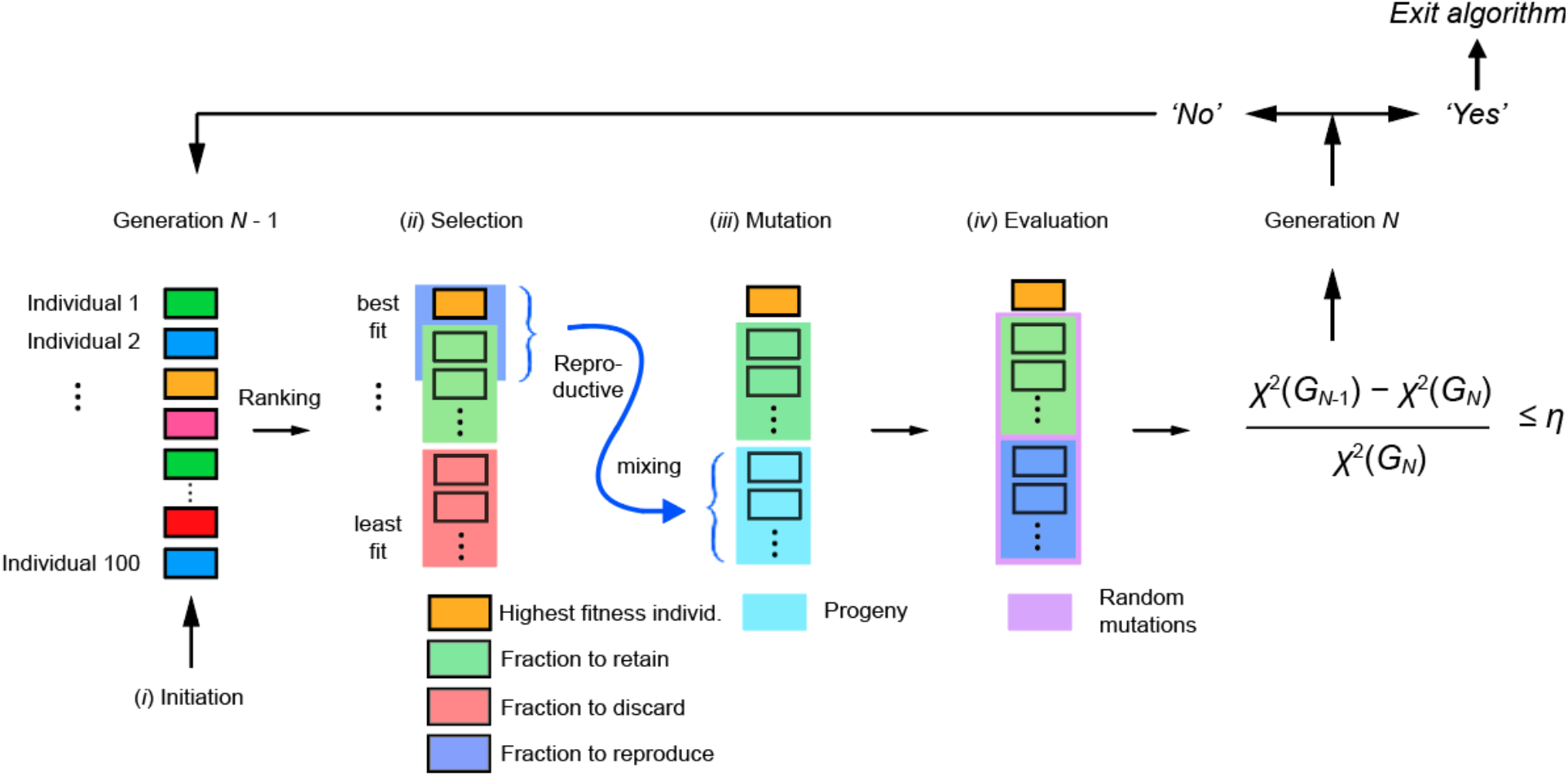
Workflow diagram of the multi-parameter optimization procedure. In order to effectively search the parameter space to obtain an optimized solution to the master equation, we applied a numerical ‘genetic algorithm’ (GA) with principles analogous to evolution by natural selection. The GA consists of four sequential steps, which are applied iteratively to successive generations of a ‘population’ of distinct solutions to the master equation. (*i*) In the ‘initialization step,’ the first-generation population is generated as a random set of trial solutions with mixed levels of fitness, as determined by the *χ*^2^ target function [see discussion of Eq. (S9)]. A population of solutions is represented in the far-left column with different colors indicating varying levels of fitness. Each ‘individual’ member of the population represents a numerical array containing a specific set of model parameters (i.e., FRET efficiency values and time constant parameters), which corresponds to a unique solution to the master equation; (*ii*) In the ‘selection step,’ the individuals are ranked according to their fitness. The fittest individuals (typically chosen as the top 50%, shaded green) are retained without alteration for the next stage of the GA, while the least fit individuals (bottom 50%, shaded red) are discarded from the population. The highest tiers of fitness are selected (typically 25%, shaded blue) for ‘reproduction,’ which involves the random exchange (or ‘mixing’) of parameters between individuals. The transformed population is represented in the third column with the reproductive fraction represented by the light-blue shaded region; (*iii*) In the ‘mutation step,’ the parameters of all but the highest-fitness individual (shaded orange) are randomly mutated to varying degrees as a means to ensure continued ‘genetic diversity.’ The final population is indicated in the far-right column with mutated individuals shaded purple; (*iv*) In the final ‘evaluation step,’ the fitness of each individual is calculated, and the one with the highest fitness is selected as the new optimized solution. If the fitness of the optimized solution is improved relative to that of the previous generation, the new generation is used as input to repeat the procedure. If the fitness of the optimized solution exceeds a preset threshold value specified by the parameter *η*, the loop is terminated. The output of the GA is then further refined using a multi-parameter linear least squares regression algorithm (MATLAB *patternsearch*) to obtain the final optimized solution.

The output parameters of the model are: (*i*) the set of time constants, 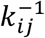 with *j* ∈ {1,2,3}, which describe the transfer times between macrostate-*i* and macrostate-*j* and (*ii*) the mean FRET efficiency values corresponding to the three macrostates, 〈*E_FRET_*〉_*i*_ with *i* ∈ {1,2,3}. We implemented a genetic algorithm (GA) to broadly search the parameter space and to obtain an approximate set of parameters that produced a relatively low value of *χ*^2^ using an arbitrary set threshold value *η*. A schematic workflow diagram for the GA optimization procedure is shown in Fig. S10. The output solution obtained from the GA was then fed into the nonlinear multivariable problem solver, *patternsearch* (MATLAB, The MathWorks, USA) to arrive at a final optimized solution. The entire procedure was repeated iteratively until the final solution was obtained. The final optimized solutions are those presented in Table S1 and Table S6.

#### Statistical Uncertainty Analysis

***We performed an analysis of the statistical uncertainty for each of the free parameters used in our model for the case of the 3’-Cy3/Cy5-oligo(dT)_15_-ss-dsDNA construct (see Fig. S11 and Table S10). This analysis is based on calculating the relative deviation of the error function*,** 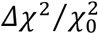**, *from the optimized value*,** 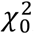**, *as a function of the time constant and FRET efficiency parameter uncertainties. The error function cross-sections given by Eq. (S9), and shown in Fig. S11, demonstrate that the optimized values we have obtained for the free parameters each correspond to a stable minimum. We assign the statistical uncertainty of these values to a 1% deviation of the error function relative to the corresponding optimized value. From this analysis, we obtain error bars for the free parameters, which we list in Table S10*.**

**Table S10.** Error bars determined from the cross-sections of the error function surfaces 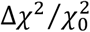, shown in Fig. S11.

| $\Delta k_{12}^{-1}$ (ms) | $\Delta k_{21}^{-1}$ (ms) | $\Delta k_{13}^{-1}$ (ms) | $\Delta k_{31}^{-1}$ (ms) | $\Delta k_{23}^{-1}$ (ms) | $\Delta \langle E_{FRET} \rangle_1$ | $\Delta \langle E_{FRET} \rangle_2$ | $\Delta \langle E_{FRET} \rangle_3$ |
| --- | --- | --- | --- | --- | --- | --- | --- |
| $\pm 0.004$ | $\pm 0.0005$ | $\pm 1.85$ | $\pm 0.03$ | $\pm 0.03$ | $\pm 0.003$ | $\pm 0.004$ | $\pm 0.005$ |

**Figure S11.**
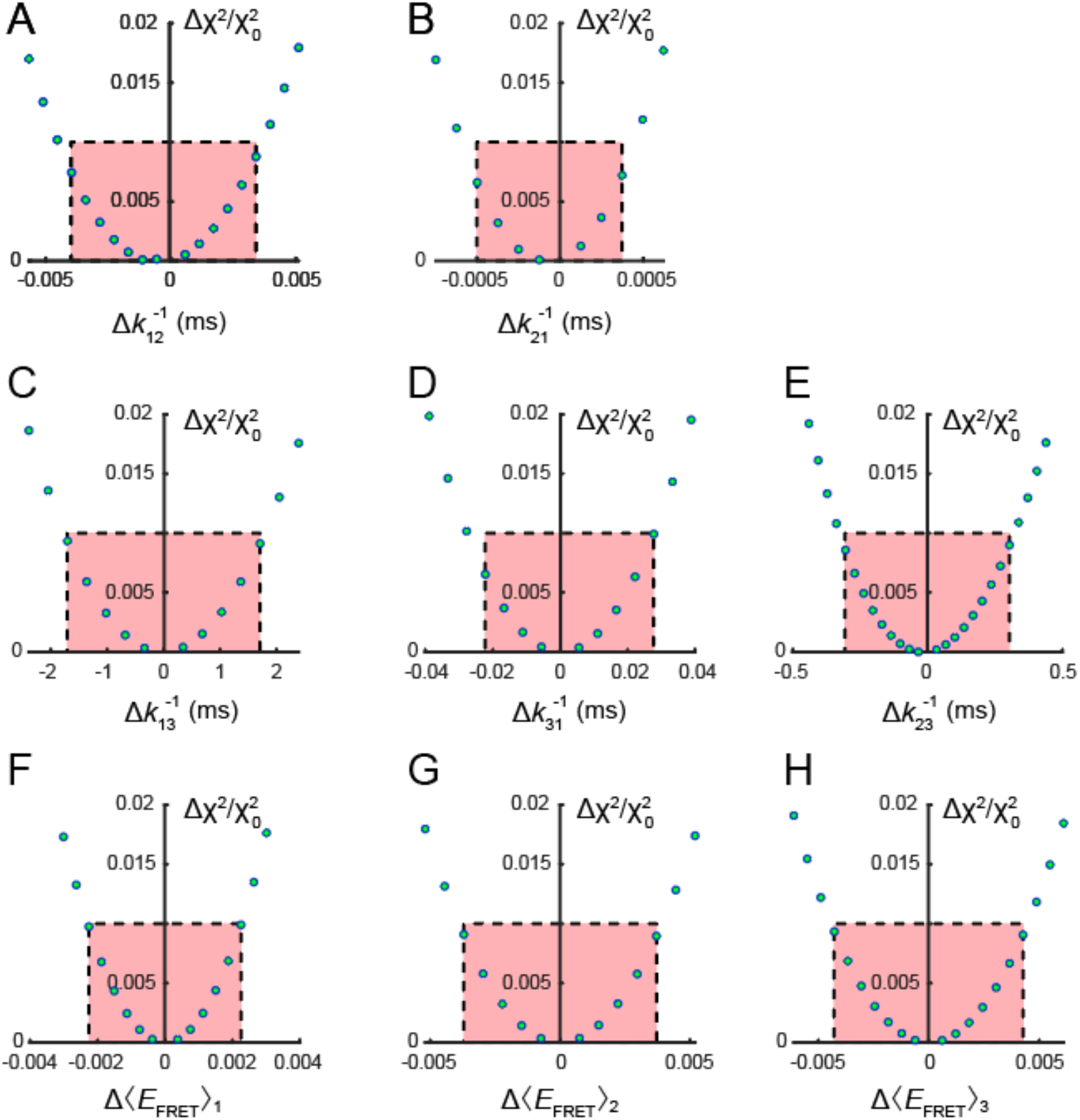
Relative deviation of the error function, 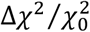, from the optimized value, 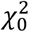, as a function of the time constant and FRET efficiency parameter uncertainties. Cross-sections of the error function given by Eq. (S9) are shown for the optimization to the 3’-Cy3/Cy5-oligo(dT)_15_-ss-dsDNA construct data set, and for the uncertainties (***A***) 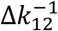, (***B***) 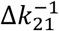, (***C***) 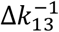, (***D***) 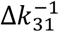, (***E***) 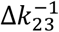, (***F***) Δ〈*E*_FRET_〉_1_, (***G***) Δ〈*E*_FRET_〉_2_, and (***H***) Δ〈*E*_FRET_〉_3_. Here Δ*x* = *x* – *x*_0_, and *x*_0_ is the value corresponding to the optimized set of parameters, which corresponds to the minimum of the multidimensional parameter surface. The red-shaded rectangles indicate the ‘confidence region’ based on a 1% relative deviation, which we associate with the quality of the experimental data. A similar analysis was performed for the data obtained for other ss-dsDNA constructs, and the cross-sections are similar to those shown above. The associated error bars reported in Table S10 are rounded up to two significant figures. We note that the time constants associated with Panels (*A*) – (*E*) are related by the detailed balance condition [Eq. (S4)], and the error bars of forward and backward elementary chemical steps were determined accordingly.

